# Novel allosteric mechanism of dual p53/MDM2 and p53/MDM4 inhibition by a small molecule

**DOI:** 10.1101/2021.12.24.474003

**Authors:** Vera V. Grinkevich, Vema Aparna, Karin Fawkner, Natalia Issaeva, Virginia Andreotti, Eleanor R. Dickinson, Elisabeth Hedström, Clemens Spinnler, Alberto Inga, Lars-Gunnar Larsson, Anders Karlén, Margareta Wilhelm, Perdita E. Barran, Andrei L. Okorokov, Galina Selivanova, Joanna E. Zawacka-Pankau

## Abstract

Restoration of the p53 tumor suppressor for personalised cancer therapy is a promising strategy. However, high-affinity MDM2 inhibitors have shown substantial side effects in clinical trials. Thus, elucidation of the molecular mechanisms of action of p53 reactivating molecules with alternative functional principle is of the utmost importance.

Here, we report a discovery of a novel allosteric mechanism of p53 reactivation through targeting the p53 N-terminus which blocks both p53/MDM2 and p53/MDM4 interactions. Using biochemical assays and molecular docking, we identified the binding site of two p53 reactivating molecules, RITA and protoporphyrin IX (PpIX). Ion-mobility mass spectrometry revealed that the binding of RITA to serine 33 and serine 37 is responsible for inducing the allosteric shift in p53, which shields the MDM2 binding residues of p53 and prevents its interactions with MDM2 and MDM4. Our results point to an alternative mechanism of blocking p53 interaction with MDM2 and MDM4 and may pave the way for the development of novel allosteric inhibitors of p53/MDM2 and p53/MDM4 interactions.

**Contribution to the field:** Given the immense importance of the p53 tumor suppressor for cancer, efforts have been made to identify p53 activators, which sterically inhibit MDM2. Because high-affinity MDM2 inhibitors are facing problems with considerable side effects, other approaches are needed to reactivate p53 for improved cancer therapy. The allosteric mechanism of action of p53 activator RITA, which we discovered, and its dependence on the oncogenic switch, is an unexpected turn in the p53 story. Our findings provide a basis for the development of p53 activators with a similar mode of functioning, either through the classical drug discovery route or through the drug repurposing approach. Allosteric modulators might have great potential as single agents, or in combination with the standard of care. Further, p53 modulators could serve as invaluable tools to better understand its biology.

## Introduction

The transcription factor p53 is a major tumor suppressor. It is a critical regulator of cell homeostasis found in multicellular organisms in the Animal kingdom (Belyi et al., 2010). p53 has a multi-domain structure and is considered an intrinsically disordered protein (IDP) (Dawson et al., 2003). p53 binds to its target promoters and regulates expression of the genes involved in cell cycle regulation and cell death. Upon activation by stress signals like DNA damage, telomere attrition, reactive oxygen species or oncogene activation, p53 is stabilized and activates or represses its target genes. p53 regulates a broad variety of cellular processes which include but are not limited to apoptosis, ferroptosis, cell cycle, DNA damage repair, senescence, metabolism, fertility or longevity (Levine, 2020).

Loss of *TP*53 alleles leads to 100% cancer penetrance in mouse models and germline *TP*53 mutations in humans predispose to the early onset of a variety of tumors (Kratz et al., 2021). The loss-of-function or gain-of-function *TP*53 gene mutations occur in approximately half of all human cancers. In tumors with intact *TP*53 gene, p53 protein is rendered functionally inert mainly due to the deregulated E3 ubiquitin ligase MDM2 and its homolog MDM4. Both MDM2 and MDM4 are amplified in cancers or undergo posttranslational modifications which promote p53 inhibition. MDM2 is an E3 ubiquitin ligase, modified by various stress signals which affect its E3 ligase activity and/or affinity to p53. Inhibition of MDM2 by stress signals increases p53 half-life and activates its transcription function (Lozano and Levine, 2016). MDM2 and MDM4 work together to direct p53 for proteasomal degradation through polyubiquitination (Vousden and Prives, 2009). MDM2 and MDM4 also inhibit p53-mediated transcription, through direct interactions with its N-terminal domain in the nucleus (reviewed in (Haupt et al., 2019)). Both proteins have p53 independent oncogenic functions and can inhibit other p53 protein family member, p73 protein (Dobbelstein and Levine, 2020).

p53 was long considered undruggable. However, based on the known crystal structure of the p53 N-terminal in complex with the MDM2 N-terminus (Kussie et al., 1996), several high-affinity inhibitors targeting the p53-binding pocket of MDM2 have been developed to date. First class of MDM2 inhibitors (MDM2i) includes nutlins, a series of cis-imidazoline analogs (IUPAC: 4-[(4S,5R)-4,5-bis(4-chlorophenyl)-2-(4-methoxy-2-propan-2-yloxyphenyl)-4,5-dihydroimidazole-1-carbonyl] piperazin-2-one) that replace the p53 α-helical peptide in MDM2 in the positions occupied by Phe^19^, Trp^23^, and Leu^26^ of p53 (Vassilev et al., 2004). The second and third class MDM2i are MI, spirooxindole compounds showing high efficacy in liposarcomas (Bill et al., 2016), RG compounds, analogs of nutlins, and piperidinones AMG-232 (Sun et al., 2014). Despite promising pre-clinical reports, idasanutlin (RG7388), the most clinically advanced MDM2i owned by Roche, failed to meet the primary end-point in Phase III clinical trial in AML and the trial was terminated due to futility (Mullard, 2020). Thus, up-to-date no MDM2i has been made clinically available yet.

Since MDM2 acts in concert with MDM4, selective MDM2i are inefficient in tumors overexpressing MDM4 protein due to structural difference in the p53 binding pocket (Toledo and Wahl, 2007; Marine et al., 2006; Jiang and Zawacka-Pankau, 2020). Several inhibitors targeting MDM4 have been developed. Yet, a great promise for improved cancer therapy lies in dual inhibitors of p53/MDM2 and p53/MDM4 inhibitors, such as α-helical p53 stapled peptidomimetic ALRN-6924, which is in phase I/II clinical development (Saleh et al., 2021).

A small molecule RITA has been identified by us in a cell-based screen for the p53 reactivating compounds (Issaeva et al., 2004). RITA restores wild-type p53 in tumor cells by preventing p53/MDM2 interaction through allosteric shift in the intrinsically disordered N-terminus of p53 and affects the binding of p53 to MDM4 (Spinnler et al., 2011; Dickinson et al., 2015). RITA has also p53-independent functions (Wanzel et al., 2016; Peuget et al., 2020).

Protoporphyrin IX (PpIX), a metabolite of aminolevulinic acid, a pro-drug approved by the FDA for photodynamic therapy topical skin lesions, reactivates p53 by inhibiting the p53/MDM2 interactions and p53/MDM4 interactions (Zawacka-Pankau et al., 2007; Jiang et al., 2019). In contrast to nutlin, neither RITA nor PpIX target MDM2, but bind to the p53 N-terminus (Zawacka-Pankau et al., 2007; Issaeva et al., 2004; Dickinson et al., 2015). However, how exactly its binding to p53 affects the p53/MDM2 and p53/MDM4 complexes remains unclear.

In the present study, we applied molecular and cell biology approaches and molecular modelling to map the region within the p53 N-terminus targeted by RITA and PpIX and to address the mechanism of their action. We found that RITA and PpIX target p53 outside of the MDM2-binding locus and identified the key structural elements in RITA molecule along with contact residues in p53, which are critical for the interaction. We found that the binding of RITA to a specific amino acid motif promotes a compact conformation of partially unstructured N-terminus, which inhibits the interaction with MDM2 and MDM4. Based on our results, we propose a model of a novel allosteric mechanism of p53 reactivation which is triggered by binding to the region spanning residues 33 – 37 of human p53.

## Materials and methods

### In situ proximity ligation assay (PLA)

*In situ* PLA was performed according to the Duolink® Proximity Ligation Assay PLA (Olink biosciences) protocol with modifications (see **Supplemental Experimental procedures** for details).

### Binding assays with [^14^C]-RITA

For a small molecule-band shift assay purified proteins (20 μM) or 80 μg of total protein from cell lysates and [^14^C]-RITA (40 μM) were incubated in buffer B (50 mM HEPES, pH 7.0, 150 mM NaCl, 35% glycerol) at 37°C/30 min and separated in standard TBE or gradient non-denaturating polyacrylamide gels. Gels were stained with Coomassie to visualize proteins and radioactivity was measured after 24-48h exposure in Fujifilm phosphor screen cassette and Phosphoimager Amersham Biosciences.

SDS-PAGE separation of cell lysates to visualize RITA/protein complexes was performed in 10% gel after snap heating at 90°C of lysates in the loading buffer. Proteins were depleted from cell lysates using anti-p53 DO-1 antibody (Santa Cruz), anti-actin antibody (AC-15, Sigma), immobilized on protein A-conjugated DynaBeads (Invitrogene).

Co-immunoprecipitation of p53/MDM2 or p53/MDM4 was performed as described previously (Issaeva et al., 2004). MDM2 in precipitates from mouse tumor cells MCIM SS cells expressing wtp53 (Magnusson et al., 1998) was detected by 4B2 antibody, a gift from Dr. S. Lain. MDMX antibody was from Bethyl laboratories.

### Scintillation Proximity Assay (SPA)

SPA PVT Protein A beads (500μl/sample, Perkin Elmer) were incubated for 2h with anti-GST antibodies (1:100). 0.1 μg/μl of studied protein in SPA buffer (GST, Np53, Np53(33/37) was added to GST-coated SPA beads. 10 μl [^14^C]-RITA (52 μCi) diluted 4 times in SPA buffer were added to protein samples (1.3 μCi). Unlabelled RITA was used as a cold substrate. SPA buffer was added to the final volume of 100 μl. Complexes were incubated for 1h at 37°C and luminescence released by the [^14^C]-RITA-excited beads was measured in standard microplate reader (Perkin Elmer).

### Circular dichroism spectroscopy (CD)

Recombinant proteins (50 μM) were incubated with RITA (reconstituted in 100% isopropyl alcohol (IPA)) or with the same amount of IPA as a blank at a 1:2 molar ratio in 25 mM ammonium acetate at 37°C for 20 min. This results in a final concentration of IPA of 5% in each case.

All CD spectra were acquired using JASCO instrument. 0.1 cm Hellma® cuvettes were used and measurements were performed in the far-UV region 260 – 195 nm at 21°C. CD spectra were recorded with a 1 nm spectral bandwidth, 0.5 nm step size with scanning speed 200 nm/min. The spectra were recorded 5 times and the data are representative of at least three independent experiments.

### Mass Spectrometry and Ion Mobility Mass Spectrometry (IM-MS)

Mass spectrometry and IM-MS were made on an in-house modified quadrupole time-of-flight mass spectrometer (Waters, Manchester, UK) containing a copper coated drift cell of length 5cm. The instrument, its operation and its use in previous studies on p53 have been described elsewhere (Jurneczko et al., 2013; Dickinson et al., 2015). Np53 was prepared at a concentration 50 μM in 50 mM Ammonium Acetate. Protein was incubated with RITA at a 1:2 molar ratio at 37°C for 30 minutes before analysis. 5% isopropyl alcohol was added to solubilize the ligand in aqueous solution, consistent with CD spectroscopy data. In all cases three repeats were taken, each on different days (For details see **Supplemental Experimental procedures**).

### Molecular Modelling

Homology model of p53 was developed using the Rosetta server (Kim et al., 2004, 2005; Rohl et al., 2004; Chivian and Baker, 2006). Generated models were validated and fitted to the cryo-EM data (Okorokov et al., 2006). Domain fitting into the 3D map of p53 was performed automatically using UCSF Chimera package from the Resource for Biocomputing, Visualization, and Informatics at the University of California, San Francisco (supported by NIH P41 RR-01081), (www.cgl.ucsf.edu/chimera/) and further refined by UROX (http://mem.ibs.fr/UROX/). (For details see **Supplemental Experimental procedures**).

### Yeast-based reporter assay

The yeast-based functional assay was conducted as previously described (Tomso et al., 2005). Briefly, the p53-dependent yeast reporter strain yLFM-PUMA containing the luciferase cDNA cloned at the *ADE2* locus and expressed under the control of PUMA promoter (Inga et al., 2002) was transfected with pTSG-p53 (Resnick and Inga, 2003), pRB-MDM2 (generously provided by Dr. R. Brachmann, University of California, Irvine, CA, USA), or pTSG-p53 S33/37 mutant and selected on double drop-out media for TRP1 and HIS3. Luciferase activity was measured 16 hrs after the shift to galactose-containing media as described previously (Inga et al., 2002; Jiang et al., 2019) and the addition of 1 μM RITA, or 10 μM nutlin (Alexis Biochemicals), or DMSO. Presented are average relative light units and the standard errors obtained from three independent experiments each containing five biological repeats. The Student’s t-test was performed for statistical analysis with p□≤□0.05-

## Results

### RITA interacts with p53 in cancer cells

Our previous findings indicate that RITA interacts with the N-terminus of p53 *in vitro* (Issaeva et al., 2004). To test whether RITA binds to p53 *in cellulo,* we analyzed the complexes of radioactively labelled [^14^C]-RITA with proteins formed in HCT 116 colon carcinoma cells carrying wild-type p53 and in their p53-null counterparts (HCT 116 *TPp53-/-*). To visualize the RITA/protein complexes, we analyzed cell lysates under mild denaturing conditions using polyacrylamide gel electrophoresis (PAGE) and detected the position of RITA and p53 by autoradiography and Western blot, respectively. Under mild denaturing conditions [^14^C]-RITA migrated with the electrophoretic front in HCT 116 *TPp53-/-* lysates (**Figure 1A**), whereas in HCT 116 cell lysates the migration was shifted, and the position of the major band coincided with that of p53. This indicates the formation of complex between p53 and RITA. Immunodepletion of p53 from the lysates using DO-1 antibody (**Figure 1A**) significantly decreased the intensity of the radioactive band supporting the band represents the p53/RITA complex.

**Figure 1.**
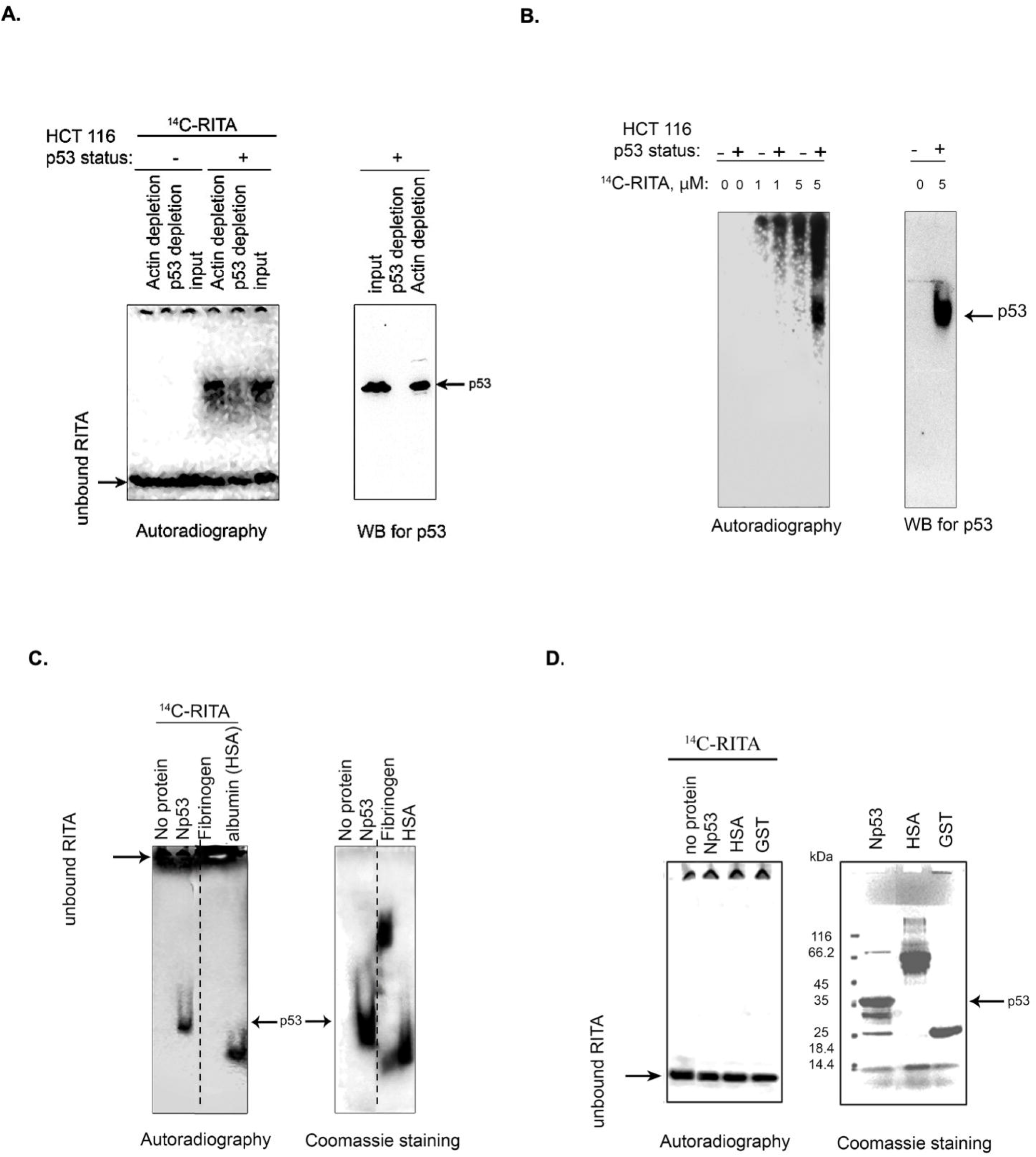
RITA binds to p53 in cancer cells and *in vitro*. **A.** [^14^C]-RITA/protein complexes were analyzed in cancer cells treated with 5 μM [^14^C]-RITA for 12h to enable sufficient accumulation of RITA. Cell lysates of HCT 116 or HCT 116 *TP53-/-* cells were separated in 10% SDS-PAGE under mild denaturing conditions (snap denaturation). The position of [^14^C]-RITA was visualized by autoradiography. p53 was detected by immunoblotting before and after immunodepletion with DO-1 antibody (right panel). Shown is a representative data of three independent experiments. **B.** A small-molecule band shift assay in gradient polyacrylamide gel run under non-denaturing conditions in TBE gel showed that [^14^C]-RITA binds to p53 in HCT 116 cells. [^14^C]-RITA and p53 were detected as in figure **A**. **C.** [^14^C]-RITA binds to GST-Np53(2-65) fusion protein, and human serum albumin (HSA) but not to fibrinogen as detected by a small-molecule band shift assay in TBE gel using 2:1 molar excess of RITA. Dotted line indicates where the gel was cut. **D.** Upon standard denaturing conditions [^14^C]-RITA/p53 and [^14^C]-RITA/HSA complexes are disrupted.

In a small-molecule band shift assay, we separated [^14^C]-RITA/cellular proteins complexes by non-denaturing electrophoresis and detected [^14^C]-RITA by autoradiography (**Figure 1B**). The major band of RITA/protein complex in HCT 116 cells coincided with that of p53 (**Figure 1B**). The absence of this radioactive band in the p53-null cells (**Figure 1B**) indicates that it represents RITA bound to p53. Taken together, our data provides evidence for the interaction of RITA with p53 in cancer cells.

### RITA interacts with the N-terminus of p53

We have shown previously that RITA interacts with N-terminal domain of p53 (Issaeva et al., 2004). Here, using small-molecule band shift assay we confirmed the interaction of RITA with the recombinant p53 N-terminus. Upon incubation, [^14^C]-RITA formed a complex with Glutathione-S-transferase (GST)-fusion p53 N-terminus (Np53) (amino acids 2-65) (**Figure 1C**) but only weakly interacted with GST-tag (**Figure 2B**). In contrast, RITA did not bind to human fibrinogen (**Figure 1C**), suggesting a selective interaction with p53. Human serum albumin (HSA), a known carrier of various drugs in blood (Koehler et al., 2002), was used as the positive drug binding control (**Figure 1C**). Under standard denaturing conditions [^14^C]-RITA/protein complexes were disrupted (**Figure 1D**), suggesting that this interaction is reversible. Non-labelled RITA readily competed out the [^14^C]-RITA from the complex with Np53 at a low molecular excess, 1:1 or 1:2.5 (Supplementary Figure S1A). However, it did not efficiently compete with the [^14^C]-RITA/HSA complex (Supplementary Figure S1B) suggesting a different mode of interaction.

**Figure 2.**
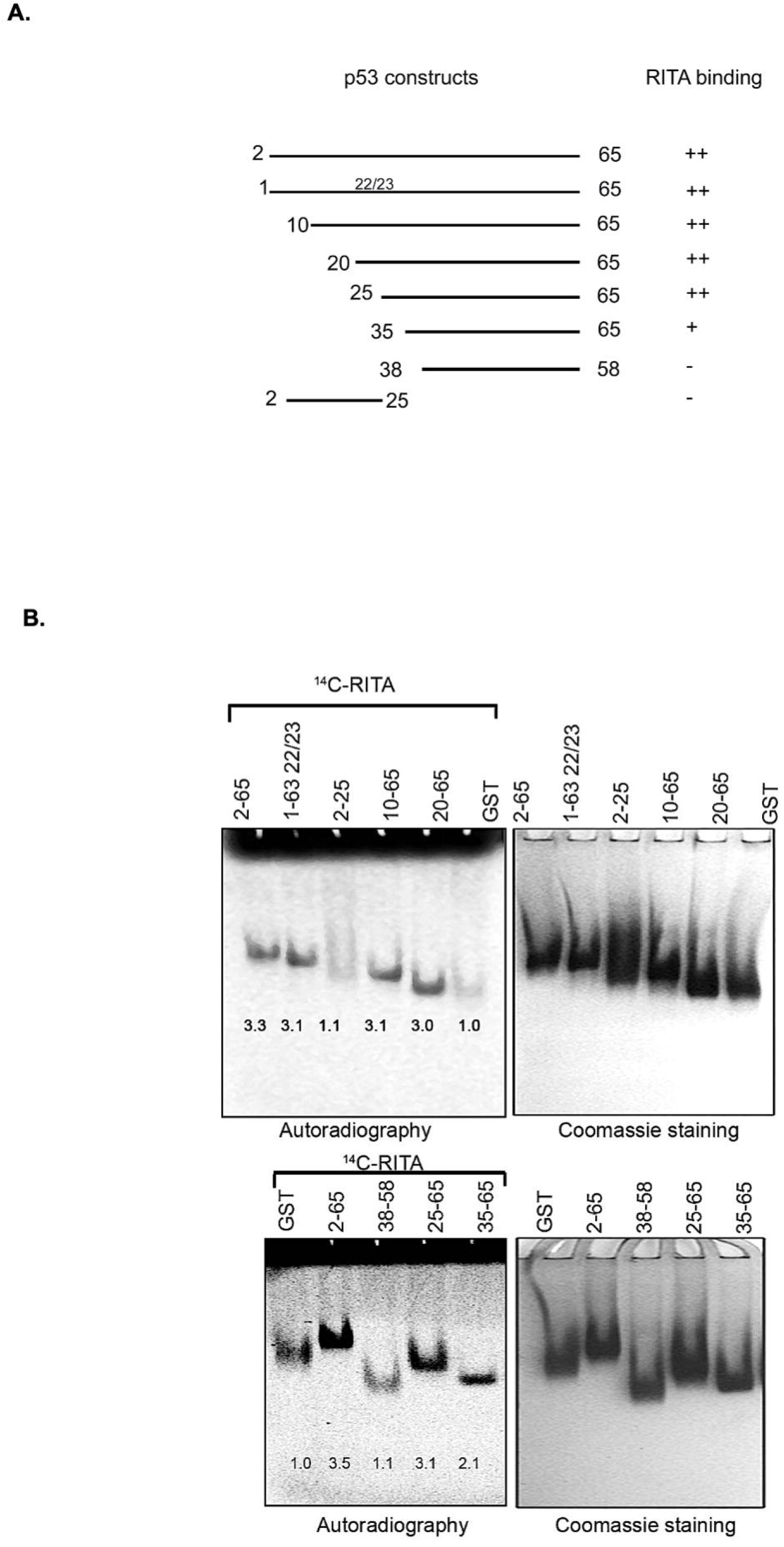
RITA binding site is located between residues 25 - 28 of the human p53 N-terminus. **A.** Scheme depicting the series of deletion mutants generated to map RITA binding site. **B.** [^14^C]-RITA only binds to p53 N terminus deletion mutants containing residues 25-38. Band density was measured using ImageJ software and normalized to GST-tag.

### RITA binding site is located in the proximity to leucine 35 in human Np53

To identify p53 residues involved in the binding to RITA, we generated a series of p53 deletion mutants and assessed their interaction with RITA (**Figure 2A**). Deletion of the first 25 residues containing the MDM2 binding site or mutations in residues 22/23 required for the interaction with MDM2 did not affect the binding of p53 to RITA, as assessed by small molecule band shift assay (**Figure 2B**, upper panel). These results argue against the binding of RITA within the MDM2 site of p53. Notably, Np53(38-58) peptide did not interact with RITA either (**Figure 2B**, lower panel). Together, our results indicate that RITA target amino acid sequence is located between residues 25-38 (**Figure 2A** and **B**). Np53(35-65) interacted with RITA approximately 50% less efficiently than Np53(2-65) (**Figure 2B**). Thus, we concluded that RITA targets residues located in the proximity to leucine 35.

### Molecular modelling reveals the binding site for RITA outside the p53 interface with MDM2

The X-ray crystallographic analysis of the p53-MDM2 complex structure shows that the N-terminal p53 region binds the MDM2 hydrophobic groove in the α-helical form (Kussie et al., 1996). The N-terminal region of p53 is largely disordered, highly flexible and forms amphipathic helical structure facilitated by the interaction with MDM2 (Wells et al., 2008). Taking into account the available information on the structural organization of the p53 N-terminus (Okorokov et al., 2006), our previous findings that RITA induces allosteric shift in p53 (Dickinson et al., 2015) and our mapping results using N-terminal mutants (**Figure 2**), we performed Monte Carlo conformational search to explore the possible binding modes of RITA to the p53 N-terminus (Schrödinger). The MCMM-LMOD search on the RITA-p53 complex found 3492 low energy binding modes within 5 kcal/mol above the global minimum. Among these, the tenth lowest energy binding mode, 2.1 kcal/mol above the global minimum, appeared reasonable with respect to the placement and orientation of RITA molecule. This model implies that the binding of RITA involves the formation of hydrogen bonds between its terminal hydroxyl groups and serine 33 and serine 37 of p53, as well as hydrophobic interactions with proline 34 and 36 via one of its thiophene and the furan rings (**Figure 3A** and **B** and **Supplemental video 1**). Hydrogen bonds and hydrophobic interactions between RITA and the p53 SPLPS amino acid sequence result in the increase of already limited flexibility of this region (**Figure 3A** and **B**).

**Figure 3.**
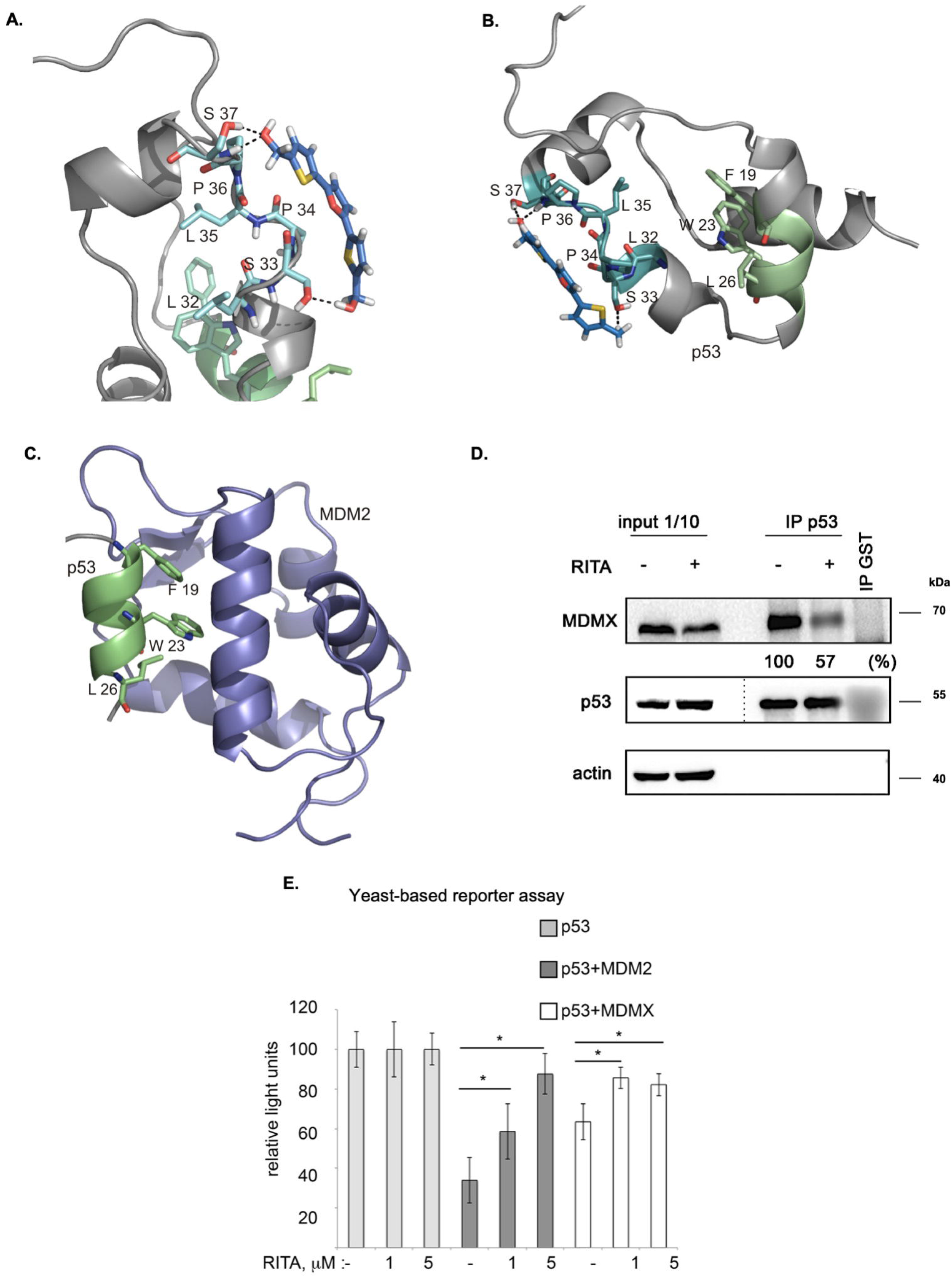
Molecular modelling shows that RITA binds to S33 and S37, induces allosteric shift in Np53 and inhibits p53/MDM2 and p53/MDM4 interactions. **A.** Binding of RITA to SPLPS sequence (cyan) of p53 involves interaction with S33 and S37 via terminal hydroxyl groups of RITA, and hydrophobic interactions with P34 and P36. Hydrogen bonds are highlighted in black dotted lines. Orientation of the MDM2-binding helix of p53 (*lime*) is different upon p53 binding to RITA (blue) **B.** and to MDM2 (*purple*) (pdb: 1YCQ) (**C**). Side chains of residues (F^19^, W^23^ and L^26^) involved in MDM2 binding are shown in (**B**, **C**.). Atom type colouring; oxygen (*red*), nitrogen (*blue*), and sulphur (*yellow*). See also Movie S1. **D.** In line with the model prediction, RITA-induced p53 conformational change results in the inhibition of p53/MDM2 and p53/MDMX binding in HCT 116 cells as assessed by co-immunoprecipitation. Dotted line indicates the site where the membrane was exposed at different exposure time. **E.** RITA rescues the p53 transcription activity from inhibition by MDM2 or MDMX as assessed by yeast-based functional assay. The average light units relative to the transactivation activity of p53 alone and the standard errors of at least five biological repeats are presented. The *t*-student test was performed for statistical analysis with p ≤ 0.05.

Molecular dynamic simulations suggest that leucine-rich hydrophobic clusters within residues 19-26 and 32-37 stabilize the folding and formation of α-helixes in the N-terminus (Espinoza-Fonseca, 2009). According to this study, the MDM2-contacting residues F^19^, W^23^ and L^26^ located in α-helix of p53 (residues 16-26) are facing inwards and are tacked inside, stabilized by the formation of hydrophobic leucine clusters, while more hydrophilic residues of the α-helix are exposed to the solvent as supported by the tryptophan fluorescence assay (Kar et al., 2002). On the other hand, the X-ray structure of the MDM2-p53 peptide complex (1YCQ.pdb) shows that MDM2-contacting residues are facing out (**Figure 3C**). This indicates that the binding to MDM2 requires a partial unwinding of the α-helix to flex out F^19^, W^23^ and L^26^, as illustrated in **Figure 3B** and **3C**. Our model indicates that RITA, by increasing the rigidity of the proline-containing SPLPS motif, induces a conformational trap in a remote MDM2 binding site. Next, we propose that constraints imposed by RITA prevent solvent exposure of F^19^, W^23^ and L^26^ residues, thus counteracting the p53/MDM2 interaction (**Figure 3B** and **3C**).

Conformational change induced by RITA is expected to impinge on other protein interactions involving the p53 N-terminus. The binding of p53 to MDM2 homolog, MDM4, requires the formation of an α-helix as well as exposure of the same p53 residues, as facilitated by MDM2. We thus reasoned that the conformational change induced by RITA might also abrogate the binding of p53 to MDM4(X).

### RITA inhibits p53/MDM4(X) interaction

Based on the allosteric shift induced by RITA in p53, we next assessed if RITA inhibits p53/MDMX complex. We treated HCT 116 colon cancer cells with RITA and assessed p53/MDMX complex inhibition by co-immunoprecipitation. Our data indicated that RITA reduced the amount of MDMX bound to p53 by 43% (**Figure 3D**).

Next, we employed a yeast-based assay, which measures the p53 transcription activity using as a readout p53-dependent luciferase reporter. Since p53 is not degraded by MDM2 in yeast cells, the inhibitory effect of MDM2 in this system is solely ascribed to the direct interaction with p53 and consequent inhibition of p53-dependent transcription (Wang et al. 2001). Co-transfection of MDM2 with p53 inhibited the p53-dependent reporter (**Figure 3E**). Notably, RITA rescued wtp53-mediated transactivation of the reporter in the presence of MDM2. Next, RITA protected p53 from the inhibition by MDMX as reflected by the restoration of p53-dependent luciferase reporter in the presence of MDMX (**Figure 3E**).

Taken together, our results demonstrated that the allosteric effects exerted by RITA result in the inhibition of both p53/MDM2 and p53/MDM4(X) interactions.

### Terminal hydroxyl groups of RITA are crucial for RITA/p53 interaction

Our model (**Figure 3**) implies that the central furan ring of RITA is not relevant for the binding with p53. Indeed, an analogue of RITA with the substitution of furan oxygen atom to sulphur (LCTA-2081, compound 2, see Supplementary Table 1 for structure) had comparable p53-dependent activity in HCT 116 cells in terms of reduction of cell viability (**Figure 4A)**. Further analysis of RITA analogues (Supplementary Table 1) showed that the presence of three rings is required for its p53-dependent biological activity. The molecular modelling predicts that one or two terminal hydroxyl groups are the key for the interaction with p53. This prediction was supported by the loss of biological activity of RITA analogue NSC-650973 (compound 4 (cpd4), Supplementary Table 1), lacking both hydroxyl groups (**Figure 4A**) and the inability of cpd 4 to compete with [^14^C]-RITA for the binding to p53 (**Figure 4B**).

**Figure 4.**
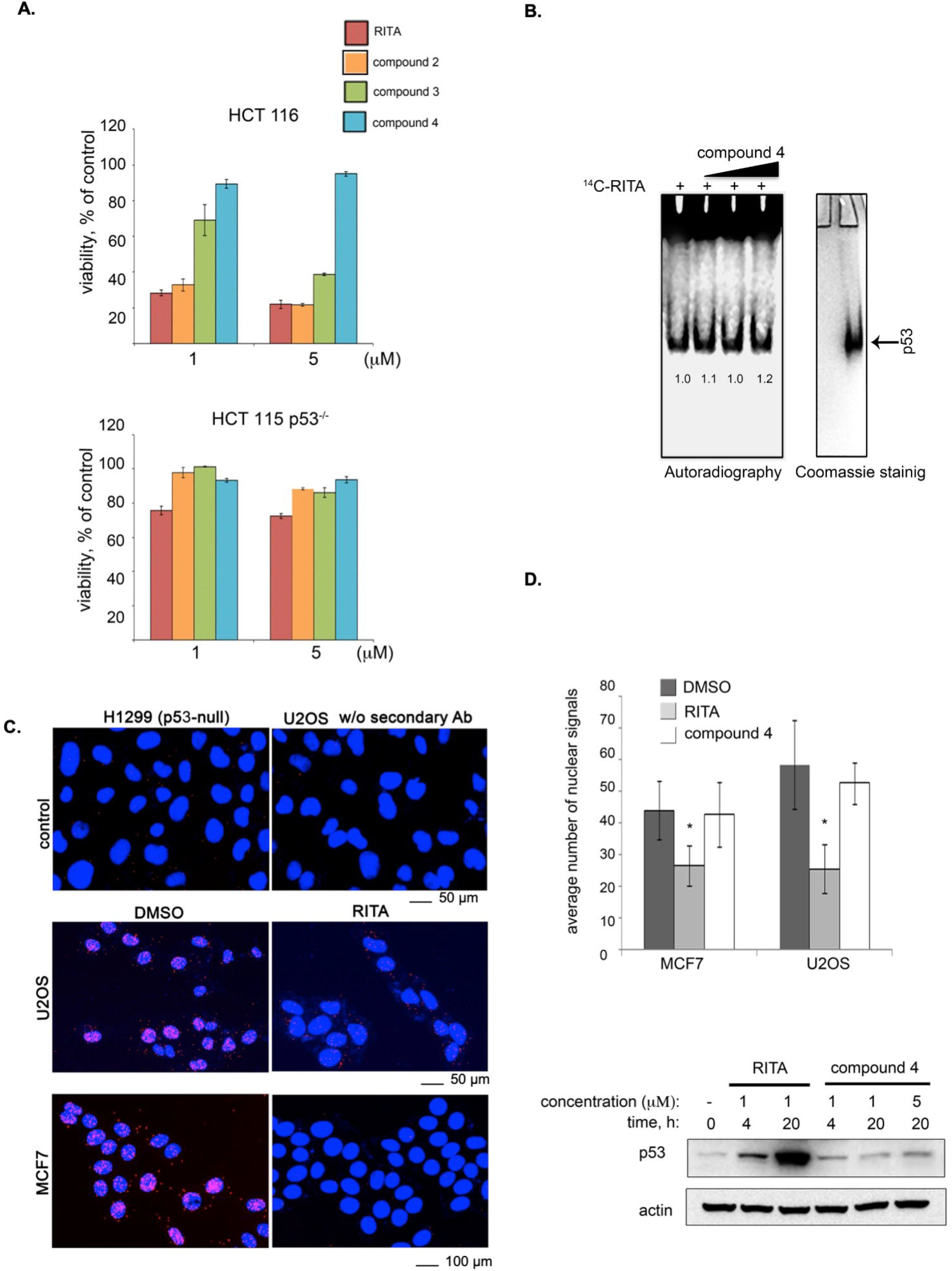
Two terminal hydroxyl groups of RITA are crucial for the binding to p53 and the inhibition of the p53/MDM2 interaction. **A.** RITA analogue NSC-650973 (compound 4) lacking two hydroxyl groups does not inhibit the growth of HCT 116 cancer cells, unlike LCTA-2081 (compound 2) analogue with substituted O atom in furan ring (for structure refer to Supplementary Table 1), which retained full biological activity. NSC-672170 (compound 3) analogue with one hydroxyl group substituted to ketone retained approximately 60% of RITA biological activity. **B.** Compound 4 (40, 80 and 100 μM) does not compete for the binding to Np53 with [^14^C]-RITA. **C.** p53/MDM2 complexes (fluorescent foci) in MCF7 and U2OS cells treated or non-treated with RITA as detected by *in situ* Proximity Ligation Assay (isPLA). The p53-null H1299 and U2OS cells stained without secondary antibody were used as the assay controls. **D.** Quantitative isPLA demonstrated the decrease in the average number of nuclear signals by RITA, but not by its derivative NSC-650973 (cpd 4) (upper panel). The normality was assessed with Shapiro-Wilk’s test. p< 0.05 values were considered statistically significant. RITA, but not compound 4 induced p53 accumulation in HCT116 cells, as detected by western blot (lower panel).

We confirmed the biological relevance of two terminal hydroxyl groups of RITA using *in situ* proximity ligation assay (isPLA) and measured the degree of inhibition of p53/MDM2 complexes in cancer cells (**Figure 4C**) (Söderberg et al., 2006; Castell et al., 2018). isPLA allows for detection of the interaction between proteins in cells using antibodies tagged to oligos. Treatment of MCF7 or U2OS cells with RITA decreased the average number of p53/MDM2 isPLA nuclei signals (from 44 +/− 9.25 to 26.5 +/− 6.45 in MCF7 cells and from 58.32 +/− 14 to 25.52 +/− 7.74 in U2OS cells when compared with DMSO) (**Figure 4D**). Unlike RITA, compound 4 did not decrease the average number of nuclei signals, indicating that it does not inhibit p53/MDM2 interaction (**Figure 4D**, upper panel). In line with these data, compound 4 did not induce p53 accumulation (**Figure 4D**, lower panel). Notably, compound 3, lacking one hydroxyl group (**Supplementary Table 1, Supplementary Figure S2B**) was more efficient in suppressing the growth of HCT 116 cells than compound 4 (NSC-650973), but still less potent than RITA **(Figure 4A** and (Issaeva et al., 2004)). Thus, we conclude that both terminal hydroxyl groups of RITA and three thiofuran rings are required for the efficient binding to p53. The ability to bind p53 correlates with the prevention of p53/MDM2 binding, induction of p53 and p53-dependent growth suppression.

### Serine 33 and serine 37 are critical for RITA/p53 interaction, p53 stabilisation and transcription activity

To further validate our model, which predicted serines 33 and 37 as RITA binding sites, we generated single and double mutant p53 proteins in which serine 33 and 37 (S33; S37) were exchanged to alanines. Next, we evaluated the binding of RITA to single and double mutants using small-molecule band-shift assay. In line with our model, the interaction of mutant p53 peptides with RITA was decreased (**Figure 5A**).

**Figure 5.**
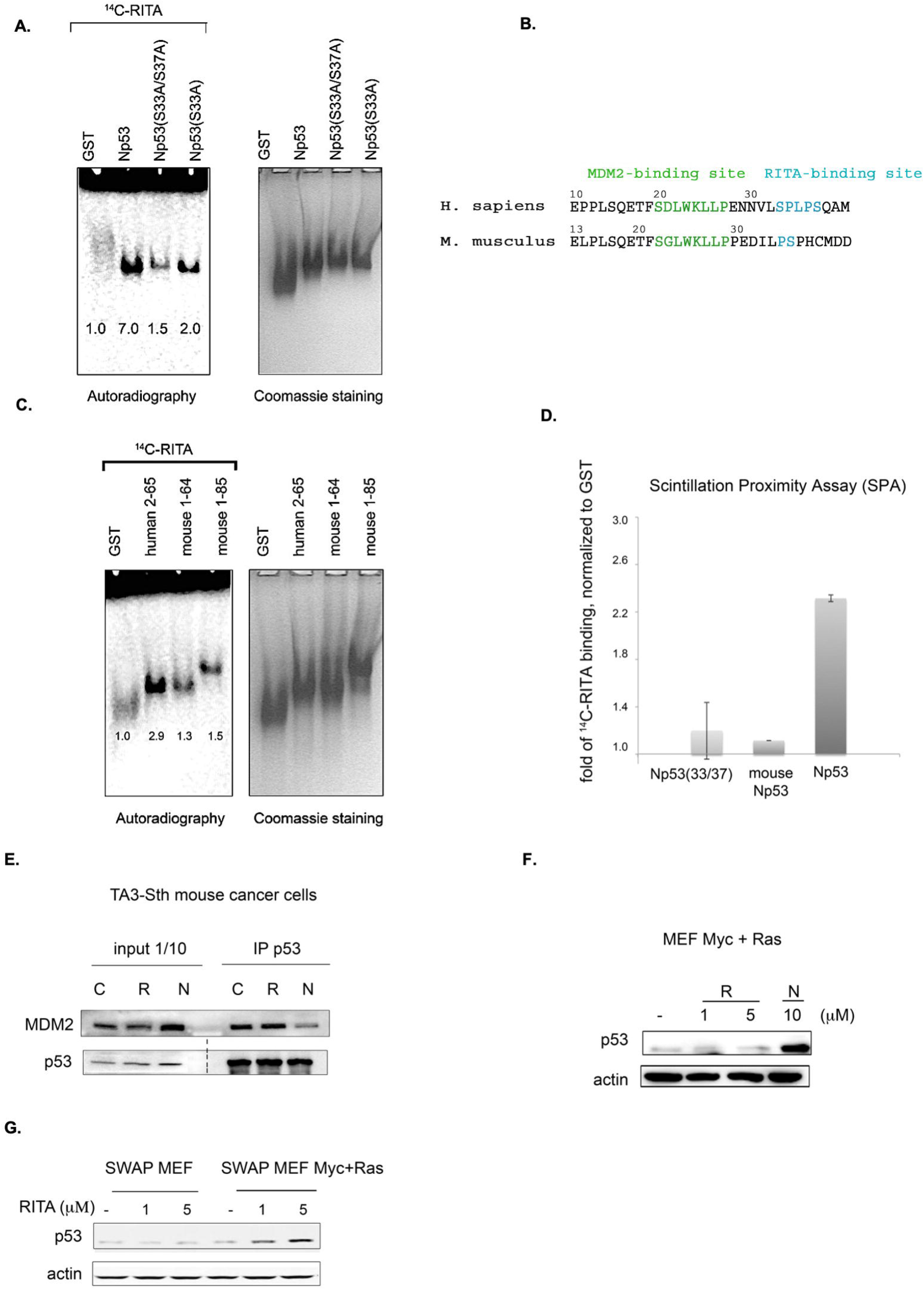
Serine 33 and serine 37 are crucial for the binding of RITA to the p53 N-terminus. **A.** Assessment of [^14^C]-RITA interaction with Np53 proteins carrying alanine substitutions of S33 or S33/S37 using band shift assay. Np53(S33/S37) does not bind to [^14^C]-RITA *in vitro* when compared to wtNp53. Bands’ densities were quantified using ImageJ software and normalized to GST-tag. **B.** Alignment of murine and human p53 N-termini. Highlighted are the sites for MDM2 interaction and the RITA-binding motif. **C. D.** RITA does not bind to mouse Np53 proteins, spanning residues 1-64 and 1-85 as detected by band shift assay and scintillation proximity assay (SPA). Np53(33/37) was used as a negative binding control in SPA assay. **D. E.** Co-immunoprecipitation showed that RITA does not prevent p53/MDM2 interaction in TA3-Sth mouse cancer cells. C - control untreated sample, R - RITA-treated, N - nutlin-treated samples; dotted line represents different exposure time of this part of the membrane. **F.** Mouse p53 in Myc- and Ras-transformed MEF’s is not induced by RITA (R) in contrast to nutlin (N). **G.** RITA induces human p53 in SWAP MEF’s transfected with Ras and c-Myc as detected by western blot.

Sequence alignment of human p53 reference protein sequence (NP_001119584.1) with murine p53 protein (NP_035770.2) showed that p53 from *Mus musculus* lacks residues corresponding to serine 33 and proline 34 (**Figure 5B**). RITA binds weakly to mouse Np53(1-64) and Np53(1-85) peptides, compared to human p53 (**Figure 5C**), suggesting that the presence of S33 and P34 is important for RITA binding. Scintillation Proximity Assay (SPA), which detects the radioactively labelled RITA only when in a very close proximity to protein coated beads showed that the binding of [^14^C]-RITA to mouse p53 and Np(33/37) mutant is inefficient in comparison with human N-terminal peptide (**Figure 5D**).

In contrast to nutlin, which blocked the p53/MDM2 complex and induced p53 accumulation in mouse cells, RITA did not disrupt mouse p53/MDM2 interaction and did not induce p53 in mouse tumor cells and mouse embryonic fibroblasts (MEFs) expressing Ras and c-Myc oncogenes (**Figure 5E** and **5F**). Nutlin but not RITA activated p53 beta-gal reporter in T22 mouse fibroblasts (Supplementary Figure S3). These data are consistent with our previous results demonstrating the absence of growth suppression by RITA in mouse tumor cell lines (Issaeva et al., 2004).

Notably, swapping mouse p53 to human p53 in mouse embryo fibroblasts (SWAP MEF) derived from transgenic mice expressing human p53 in mouse p53-null background (Dudgeon et al., 2006) restored the ability of RITA to reactivate p53. RITA induced p53 in SWAP MEF’s expressing c-Myc and Ras (**Figure 5G)**. It did not affect the viability of SWAP cells without Ras and Myc overexpression, which is in line with our previous data suggesting that oncogene activation is required for RITA-mediated induction of p53 (Issaeva et al., 2004; Grinkevich et al., 2009). Taken together, these data suggest that S33 within SPLPS motif is required for RITA binding to p53.

Next, we compared the ability of RITA to rescue the transcriptional activity of p53 and S33A/S37A mutant from MDM2 using yeast-based reporter assay. Both nutlin and RITA prevented MDM2-mediated inhibition of p53 activity (**Figure 6A**). However, RITA did not rescue from MDM2 the transcriptional activity of p53(33/37), while nutlin protected both wt and p53(33/37) from inhibition by MDM2 (*t*-student; p < 0.05) (**Figure 6A**). These data support the notion that S33 and S37 play an important role in RITA-mediated inhibition of p53/MDM2 interaction.

**Figure 6.**
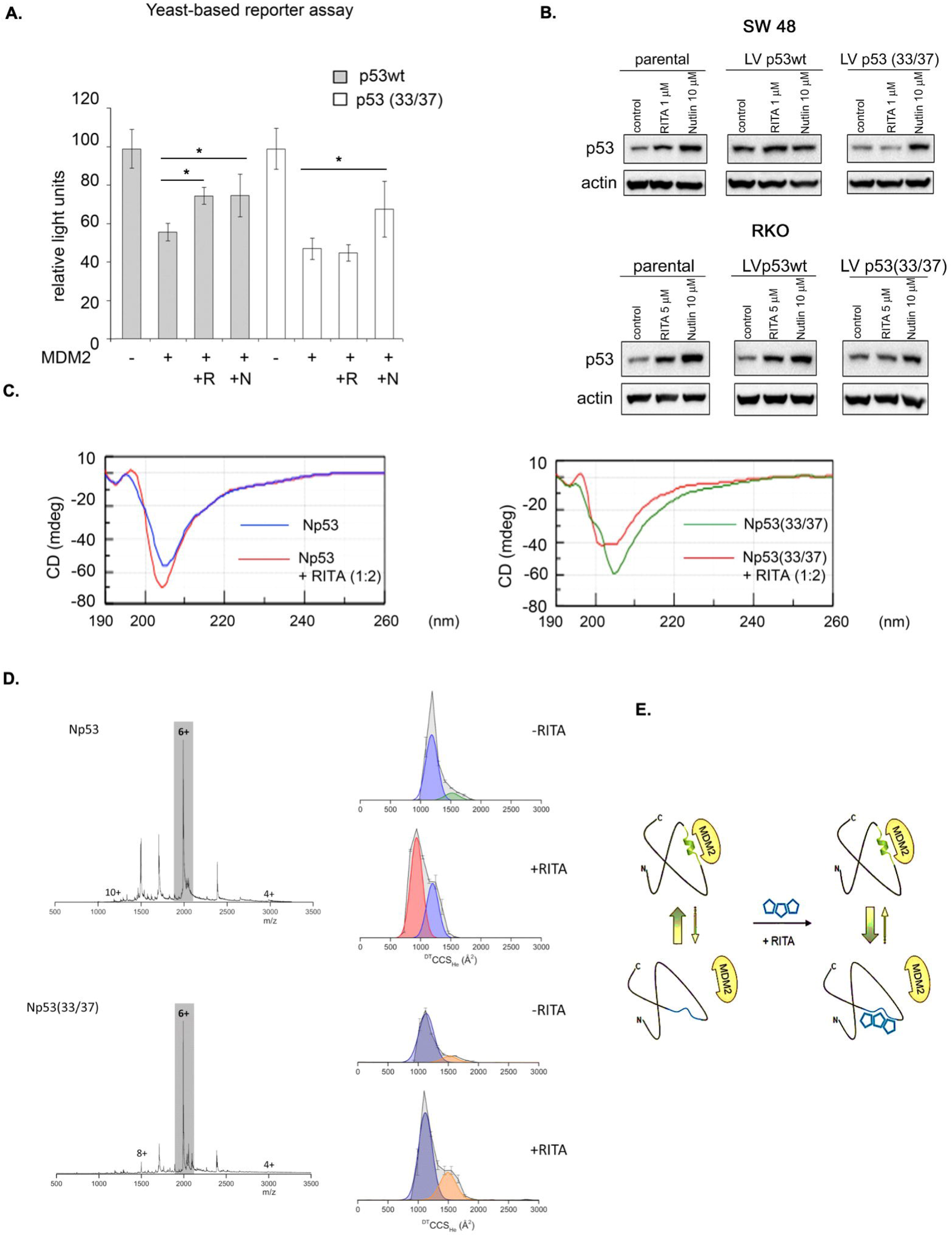
Serine 33 and serine 37 are required for RITA-induced rescue of p53 from inhibition by MDM2. **A.** Co-expression of MDM2 along with wt or mutant p53(33/37) inhibits p53-dependent luciferase reporter in yeast-based reporter assay. 1 μM RITA (R) does not rescue the reporter driven by mutant p53(33/37). The *t*-student test was performed for statistical analysis with p<0.05. (N) - nutlin **B.** wt p53 and mutant p53(33/37) were overexpressed using lentivirus in RKO *TP*53-/- and SW 48 cancer cells. Wt p53 protein but not mutant p53(33/37) is induced by 1 μM RITA (R) as assessed by western blotting. Band density was assessed using ImageJ software and normalized to non-treated controls. (N) - nutlin **C.** RITA increases the secondary structure content in wt Np53 (left) but not in mutant Np53(33/37) (right) as detected by circular dichroism spectroscopy (CD). **D.** nESI mass spectra (left) and drift tube ion mobility mass spectrometry collision cross section distributions arising from arrival time distributions (right) for the [M+6H]^6+^ analyte of wt Np53 in the absence and presence of RITA (top panel, and mutant Np53(33/37) in the absence and presence of RITA (bottom panel). Conformational families are depicted by coloured Gausian curves. wtNp53 undergoes a compaction event resulting in the induction of a novel conformational family shown in red. Mutant Np53(33/37) conformational spread is unaffected by RITA. **E.** A scheme illustrating allosteric mechanism of RITA-induced inhibition of p53/MDM2 interaction. Binding of RITA shifts the balance towards p53 conformation with low affinity to MDM2.

To assess if serine residues are important for the induction of p53 in human cells by RITA, we overexpressed S33/S37 p53 and wtp53 in colon carcinoma RKO *TP53-/-* cancer cells. Nutlin induced the accumulation of wt and p53(33/37) with similar efficiency (**Figure 6B** and not shown). In contrast, the induction of the double serine mutant by RITA was impaired (**Figure 6B** and not shown).

CD spectroscopy confirmed our previously published data (Dickinson et al. 2015) that RITA increases the content of the secondary structure in Np53 (**Figure 6C,** left panel). Thus, RITA binds to S33, S37 and induces a conformational change in p53 that inhibits p53/MDM2 complex and induces p53 stabilisation (**Figure 6C**, left panel). To elucidate the role of serine 33 and 37 in RITA-mediated increase of the secondary structure in Np53, we incubated wtNp53 and Np53(33/37) with the access of RITA (1:2 ratio) and performed CD measurements. As shown in **Figure 6C**, (right panel), RITA did not increase the secondary structure content in Np53(33/37) when compared to wtNp53. The induction of the allosteric shift in wtNp53 and in Np53(33/37) by RITA was next analyzed by ion mobility mass spectrometry (IM-MS) that was described by us previously (Jurneczko et al., 2013; Dickinson et al., 2015). Briefly, we first incubated both wt and Np53(33/37) in the presence or absence of RITA (Supplemental Experimental procedures). The wtNp53 after incubation with RITA presents as ions of the form [M+zH]^z+^ where 4≤ *z* ≤10 with charge states 5≤ *z* ≤ 8 at significant intensity (**Figure 6D,** upper panel). The mass spectra for Np53 without RITA, with RITA and the control spectra show no mass shift, suggesting that RITA binding is lost during desolvation (Supplementary Figure S4A). Thus, RITA changed the conformation of Np53 as described previously (Dickinson et al., 2015). The collision cross section distributions in **Figure 6D,** show that in the absence of RITA, Np53 presents in two distinct conformational families centered at ~1500 and ~1750 Å^2^. After incubation with RITA the more extended conformer is lost, the conformer at ~1500 Å^2^ remains present at a lowered intensity and a third conformational family centered on ~1000 Å^2^ appeared, suggesting a significant compaction of the Np53 protein. Control experiments confirmed that this conformer was present only after incubation with RITA (Supplementary Figure S4A). This trend was observed for all sampled charge states (Supplementary Figure S5A) with small variations in conformer intensity attributable to coulombic repulsion upon desolvation. Thus, RITA induces a unique compact conformer (or closely related conformational family) in wtNp53. In contrast, no gross conformation change was detected in mutant Np53(33/37) after incubation with RITA (**Figure 6D**, bottom panel and Supplementary Figure S4B, S5B). Np53(33/37) was present in two conformations centered at ~1100 and 1500 Å^2^ both in the absence and presence of RITA.

In summary, using a number of experimental approaches, we identify p53 residues S33 and S37 crucial for the interaction with RITA and the allosteric activation of p53.

### PpIX is an allosteric activator of p53

Next, we assessed whether the allosteric mechanism of p53 activation identified by us applies to other inhibitors of p53/MDM2 interactions. Through drug repurposing approach, we have previously shown that small molecule protoporphyrin IX (PpIX), a drug approved to treat actinic keratosis, binds to the p53 N-terminus and disrupts p53/MDM2 and p53/MDMX complexes (Zawacka-Pankau et al., 2007; Sznarkowska et al., 2011; Jiang et al., 2019). Here, we tested if PpIX targets the same amino acid residues in p53 as RITA, using fluorescent-based small-molecule band shift assay. Fluorescent band shift assay indicates that substitution of serine 33 to alanine or double substitution at serine 33 and serine 37 decreases the binding of PpIX to the p53 N-terminus (**Supplementary Figure S6**).

Taken together, our findings implicate the conformational state of the SPLPS sequence distal from the MDM2-interacting residues as a key structural element regulating p53/MDM2 interaction as presented in model in **Figure 6E**.

We propose that this site could be modulated by small molecules such as RITA and PpIX to reactivate p53 for improved cancer therapy.

## Discussion

Reconstitution of the p53 tumor suppressor has proven to induce regression of highly malignant lesions (Junttila et al., 2010) and several compounds targeting the p53/MDM2 interaction *via* steric hindrance are currently undergoing clinical trials (Jiang and Zawacka-Pankau, 2020). Yet, unexpected toxicities observed in clinical studies demand the identification of novel compounds with a distinct mode of action.

RITA reactivates wild-type p53 and inhibits p53/MDM2 interaction, however, it is unique among known p53/MDM2 inhibitors because it binds to p53 (Issaeva et al., 2004; Dickinson et al., 2015). Even though RITA has been reported to display p53-independent functions (Wanzel et al., 2016; Peuget et al., 2020), it is a valuable tool to explore the mechanism of wild type p53 reactivation.

The gel shift assays employed by us demonstrated that RITA binds to p53 in cells and *in vitro.* Mutation analysis and molecular modelling identified S33 and S37 as critical residues responsible for RITA binding to Np53. Importantly, the binding of RITA inhibits both p53/MDM2 and p53/MDMX interactions (**Figure 1 & Figure 2, 3**).

Molecular dynamic simulations showed that residues 32-37, responsible for RITA binding, might be involved in the stabilization of a conformational state in which MDM2-contacting residues F^19^, W^23^ and L^26^ of p53 are tacked inside the molecule. Since X-ray structure of the MDM2 in complex with short p53 peptide (1YCQ.pdb) suggests that residues F^19^, W^23^ and L^26^ should be facing out in order to bind MDM2, as shown in **Figure 3C**, thus the binding of p53 to MDM2 requires conformational changes. More recent study revealed that segments 23-31 and 31-53 of the p53 N-terminus are involved in long-range interactions and can affect p53’s structural flexibility upon MDM2 binding or phosphorylation of residues S33, S46 and T81. In particular, non-random structural fluctuations at 31-53 segment are affected by MDM2 binding (Lum et al., 2012). These data provide important evidence supporting our idea that restricting conformational mobility of segment involving residues 33-37 might serve to prevent the p53/MDM2 interaction.

Our previously published data with ion-mobility mass spectrometry (IM-MS) (Dickinson et al., 2015) and molecular modelling (**Figure 2**) implies that RITA binds weakly to p53 and induces allosteric shift in Np53. Modulation of protein conformation by a weak binding ligand has previously been shown by IM-MS (Harvey et al., 2012). Since IM-MS detects the changes in p53 conformation induced by a single point mutations (Jurneczko et al., 2013), analogous to the structural changes in Np53 induced by RITA as detected by IM-MS, we analyzed the conformer states of the double mutant Np53(33/37) (**Figure 6**). The substitution of S33 and S37 to alanines and incubation with RITA do not significantly affect mutant Np53 conformers when compared to wt Np53. We confirmed that the finding that RITA does not change the conformation of the double mutant Np53(33/37) using CD spectroscopy. Next, RITA inhibited p53/MDM2 and p53/MDMX complexes in yeast-based reporter and in cancer cells (**Figure 2 & 3 & 4**). Yet, the MDM2 and MDMX complexes with double mutant Np53(33/37) were only inhibited by nutlin but not by RITA. Thus, S33 and S37 are crucial for allosteric shift in Np53 induced by RITA.

Sequence alignment analysis showed that murine p53 lacks S33 and P34. Functional analysis revealed that RITA does not bind to mouse Np53 *in vitro* and does not reactivate wt p53 in murine cancer cells (**Figure 5**), which further supports the significance of S33 and S37 in p53 reactivation by RITA.

Next, using drug repurposing, we found that PpIX, which we have shown to bind to p53 and to disrupt p53/MDM2 and p53/MDMX interaction (Jiang et al., 2019), also requires serine 33 and 37 for p53 reactivation.

Allosteric mechanism of p53 reactivation by RITA and PpIX is a novel and promising turn in the development of inhibitors of p53/MDM2 interaction. Recent studies showed that epigallocatechin-3-gallate (EGCG) binds to ITD Np53, inhibits p53/MDM2 interactions and reactivates p53 by preventing proteasomal degradation (Zhao et al., 2021) highlighting the relevance of our discovery.

**Figure 6E** illustrates a scenario suggested by us, in which p53 exists in a range of conformational states, that are present in cells in a dynamic equilibrium. Close proximity to MDM2 induces F^19^, W^23^ and L^26^ to be exposed and to fit into the p53-binding cleft of MDM2, causing the equilibrium to shift in favour of this conformation. Binding of RITA and PpIX to SPLPS stabilizes the alternative conformation, in which MDM2-contacting residues are trapped inside. In this way, the binding of RITA and PpIX to p53 shifts the balance towards the p53 conformer with low affinity to MDM2 and likely to MDMX.

In summary, our data establish that the allosteric mechanism of inhibition of p53/MDM2 and p53/MDMX interaction by small molecules could be a viable strategy for the development of p53-reactivating therapies with the mode of action different from MDM2i. The identified structural elements in p53 and RITA may provide a basis for the generation of novel allosteric activators of p53, which might be translated into the clinical practice in a future.

## Supporting information

Supplemental material description

## Supplemental information

Supplemental information includes Supplemental Experimental Procedures, Supplemental References, six supplemental figures and one table.

## Acknowledgements

This work was supported by grants to G.S. from the Swedish Research Council, the Swedish Cancer Society and Ragnar Söderberg Foundation. ALO was funded by BBSRC UK. J. E. Z-P would like to acknowledge the grant from Karolinska Institute, Stockholms Läns Landsting, the Strategic Research Program in Cancer Karolinska Institute, Åke Wibergs Stiftelse and Cathrine Everts forskningsstiftelse. J E. Z-P would like to address special thanks to Klas Wiman from Karolinska Institute for his support and outstanding mentorship. We are greatly indebted to Protein Science Facility Karolinska Institutet for protein purification and to all our colleagues who shared with us their reagents and cell lines. The authors are grateful to Yari Ciribilli, Bartosz Ferens, Anna Kostecka and Alicja Sznarkowska for helpful discussions and technical assistance.

## Authors contribution

Conceptualisation: G. Selivanova; J. E. Zawacka-Pankau; A.L. Okorokov

Methodology and data analysis: G. Selivanova; J. E. Zawacka-Pankau; A.L. Okorokov; V.V.

Grinkevich; N. Issaeva; P.E. Barran; A. Inga; LG. Larsson; A. Karlen

Investigation: J.E. Zawacka-Pankau, V.V. Grinkevich, A.Vema, K. Fawkner, N. Issaeva, V.

Andreotti, A. Inga, E.R. Dickinson, E. Hedström, C. Spinnler, M.Wilhelm

Writing draft: J. E. Zawacka-Pankau, A.L. Okorokov, A. Inga, P.E. Barran, G. Selivanova

Writing review and editing: J. E. Zawacka-Pankau, G. Selivanova;

Supervision: G. Selivanova and J. E. Zawacka-Pankau.

