## Supplemental material description for "Novel allosteric mechanism of dual p53/MDM2 and p53/MDM4 inhibition by a small molecule"

**Supplemental methods**

**Chemical compounds**

RITA (2,5-Bis(5-hydroxy[^14^C]methyl-2-thienyl) furan, NSC652287), NSC 650973-N, NSC 672170 and [^14^C]-RITA (19 mCi/mmol) was obtained from Research Triangle Institute, purchased from Tjaden Biosciences, or synthesized by us (LCTA-2081). Nutlin, and protoporphyrin IX were purchased from Sigma-Aldrich.

**Drug treatments**

When not indicated RITA was used at 1 μM concentration and nutlin3a at 10 μM. Protoporphyrin IX was used at 5 μg/ml.

**Protein expression and purification**

GST-fusion proteins were purified as described (Issaeva et al., 2004; Okorokov et al., 2006). Constructs encoding deletion mutants of GST-Np53 were obtained by cloning corresponding PCR fragments into the pGEX-2T vector. Alanine substitution of residues 33 and 37 was introduced using site-directed mutagenesis kit (Stratagene). Constructs encoding mouse GST-p53 (1-64) and (1-85) were a generous gift from Dr. D. Meek. Dr. R Iggo kindly provided lentivirus constructs for the expression of wild-type and (33/37) mutant p53 in cancer cells. Np53 and Np53(33/37) peptides without a tag for CD and IM-MS studies were purified by Protein Science Facility, Department of Medical Biochemistry and Biophysics, Karolinska Institutet.

**Cell-based assays**

Series of colon carcinoma cell lines with deleted p53 (HCT 116 *TP53-/-*, RKO *TP53-/-*, SW 40 *TP53-/-*) as well as parental wtp53 expressing HCT 116, RKO cells were kindly provided by B. Vogelstein.

Swap MEFs were a gift from E. Appella. TA3-Sth mouse mammary gland carcinoma, U2OS human osteosarcoma, MCF7 human breast cancer, H1299 human lung adenocarcinoma and were obtained from ATCC. Cell viability was determined in short-term viability assay using WST-1 (Roche) proliferation reagent according to the manufacturer protocol.

**In situ proximity ligation assay (PLA)**

Cells were grown in 8-well chamber slides (VWR), treated with 5 μM MG132 for 30 min before treatment with 5 μM RITA/6 hours. After washing with cold PBS the cells were fixed in 70% ethanol at -20°C and blocked using 40 μl of blocking buffer (3% BSA, 2.5 ng/ml ssDNA, 2.5mM L-Cysteine, 50 mg/ml RNase A, 5mM EDTA, 0.05% Tween 20 (all from Sigma) in PBS) at 37°C/2 hours. Rabbit anti-p53 antibody FL393 and mouse anti-MDM2 SMP14 (both Santa Cruz Biotechnology) at 1:50 dilution in blocking buffer without RNAse A were applied overnight at 4°C. After washing 3x2 min with TBS/0.1%, Tween 20 the PLA probes (a–mouse MINUS and a–rabbit PLUS) were applied (1:10 dilution), along with DAPI stain (Sigma) and incubated for 1 hour at 37°C. Detection Kit 563 was used according to the manufacturer’s protocol (Olink biosciences). Analysis and picture acquisition were performed using a Leica DMRE fluorescence microscope with a Hamamatsu dual mode cooled CCD C4880 digital camera and the HiPic software. The number of fluorescent dots per nuclei (>100 nuclei per sample form three independent experiments) was calculated using BlobFinder Imaging software (<http://www.cb.uu.se/~amin/BlobFinder>). The Student’s *t*-test was performed for statistical analysis with p ≤ 0.05.

**Ion mobility mass spectrometry**

Ions were produced by positive nESI with a spray voltage of 1.35-1.6 kV. IM-MS experiments were performed with helium as the buffer gas; its pressure measured using a baratron (MKS Instruments, UK). The drift voltage across the cell was varied by decreasing the cell body potential from 60 V to 15 V, with arrival time measurements taken at a minimum of five distinct voltages. Temperature and pressure readings were taken for each drift voltage and used in the analysis of drift time measurements (approximately 28°C and 3.5 Torr, respectively). nESI tips were prepared in-house with a micropipette puller (Fleming/Brown model P-97, Sutter Instruments Co., USA) using 4” 1.2 mm thin wall glass capillaries (World Precision Instruments, Inc., USA) and filled with 10-20 μL of the sample at the specified concentration. Tuning conditions during the experiments were kept as constant as possible: cone voltage 65-70 V, source temperature 80°c, pusher period 104-112 µs.

Data were analyzed using MassLynx v4.1 software (Waters, Manchester, UK), Origin v8.5 (OriginLab Corporation, USA) and Microsoft Excel. Ion arrival time distributions were recorded by synchronization of the release of ions into the drift cell with the mass spectral acquisition. The CCS distribution plots are derived from raw arrival time data at a drift voltage of 35 V using Equation 1 (Mason and McDaniel, 1988; McCullough et al., 2008).

$\Omega_{\mathrm{avg}} = \frac{\left( 18\pi\right)^{1/2}}{16}\left[ \frac{1}{m_{b}}+ \frac{1}{m} \right]^{1/2}\frac{ze}{\left( K_{B}T \right)^{1/2}}\frac{1}{\rho}\frac{t_{d}V}{L^{2}}$	Equation 1

Where m and m_b_ are the masses of the ion and buffer gas, respectively; z is the ion charge state; e is the elementary charge; K_B_ is the Boltzmann constant; T is the gas temperature; ρ is the buffer gas density; L is the drift tube length; V is the voltage across the drift tube; and t_d_ is the drift time.

**Molecular modeling**

The best-fitted model was used for the docking studies. The initial structure of the p53-RITA complex was generated by manually placing RITA close to the experimentally identified binding site (residues 31-38) in the Cryo-EM fitted model. The generated complex was minimized using the OPLS-2005 force field with “normal” BatchMin cutoffs (7.0Å VDW; 12.0Å ELE). Possible binding modes of RITA were explored using 10000 steps of Monte Carlo conformational search implemented in Mixed torsional/Large Scale Low mode sampling algorithm (MCMM-LLMOD) (Kolossváry and Guida, 1992) within MacroModel (MacroModel, version 9.6; Schrödinger). The binding site residues (31-38) were allowed to move freely, while the remaining residues were treated as frozen. The MOLS command in BatchMin was used for the simultaneous translation/rotation of the ligand by random variations between 0-3 Å and 0-180º, respectively. Low energy protein-ligand complexes found within 5 kcal/mol above the global minimum were saved for visual inspection.

**Generation of mouse embryonic fibroblasts transformed with Myc and Ras**

Cells were transfected with pVEJBRas and PSPmyc vectors using Lipofectamine 2000 (Invitrogen), stable clones were selected in the presence of the selective antibiotic.

Western Blot was done according to standard procedure, membranes were incubated with primary rabbit p/c p53 FL-393 (Santa Cruz), rabbit p/c Myc N-262 (Santa Cruz), rabbit p/c Ras 3965 (Cell Signalling), mouse m/c b-actin (Sigma) and secondary goat anti-mouse IRDye 680 and goat anti-rabbit IR Dye 800 (Licor), and developed using Li-Cor Odyssey Imaging System.

**Βeta-gal p53 reporter assay**

ARN8 (human melanoma) and T22 (mouse fibroblasts) RGC-ΔFos-*lacZ* cell lines, which are expressing lacZ- gene under p53-dependent promoter, were a nice gift from Dr. S. Lain, Karolinska Institutet. The assay was performed as described previously (Berkson et al., 2005; Lain et al., 2008). Briefly, cells were seeded in 96-well plate at the desired density and treated with 10 μM Nutlin or 1 μM RITA for 24 h. Following incubation, the activation of p53 is detected by using the CPRG mix solution. The absorbance was detected in a microplate reader PerkinElmer, at 570 nm after 24 h incubation.

**Details of molecular docking experiments.**

Rules for chemical structures defragmentation:

- the following bond types - C-N, N-N, N-O, C-C, S-S, C-O as well as bonds between the ring atom and non-ring atom and bonds connecting two cycles can be split; double and triple bonds cannot be broken.

Molecular docking was performed using Autodock 4.0 using Compute Cluster Pack for the parallel computational procedures. The calculations were implemented using cluster HP (33x2xAMD Opteron 2.6 GHz double-core, 16x12GB+16 RAM, 24x73GB HDD, Infinity Band (MPI interconnect), Gigabit Ethernet, Windows 2003 Compute Cluster edition). Computational docking procedure allows obtaining about 50000 virtual complexes for one macromolecule target per 24 hours with maximal utilization of computer facilities. Crystal structure of p53 has not been resolved. We used the model of p53, which was fitted into cryoelectron microscopy imaging of purified ATP-stabilised murine p53 protein. The model of p53 tetramer was implemented by Andrei L. Okorokov (Okorokov et al., 2006). The resolution of the model is 13.7 A°. Since it is suggested that mdm2-p53 interaction site is different from the site of the p53-tetramerization domain (Moll and Petrenko, 2003), we used in calculations one of the four identical subunits.
To study the behavior of p53 model at the normal conditions, molecular dynamics was performed (Amber force fields, 300K, 1 ns). The results of the dynamics demonstrated the stability of the model at normal conditions during 1 ns.

The grid’s elements in Autodock4.0 are formed a cube. To investigate the mode of RITA binding close to SPLPS motif, the center of was positioned near the peptide 30-40, and the distance between the center of the cube and the plane of each surface was 14.3 Ǻ.
Initial conformation of every small-molecule structure was defined using Concord Standalone (part of Sybyl8.1) which allows converting the 2D structure into a 3D structure.

The initial position of small molecules in the grid was selected randomly without any suggestion about the location of the molecule in the N-terminal of p53.

To search for the global minimum of protein-ligand binding energy, we used genetic algorithm available in Autodock 4.0. The genetic algorithm is based on the principle of natural selection. During the simulation (modeling) process some generations with a specific initial conformations of protein and ligand (number of generations) are produced. The number of generations equals 27,000. Change of conformations and evaluation of energy is estimated for each generation (number of individuals). We defined the number of individuals as 100. By calculated binding energy, several conformations can be selected for the next generation (number of individuals selected according to the fitness function). We set 1 as a value for this parameter of the genetic algorithm.

The Estimated Free Energy of Binding (*E_bind*) in Autodock 4.0 consists of four components:

*E_bind = Final Intermolecular Energy + Final Total Internal Energy + Torsional Free Energy – Unbound System energy.*

*Final Intermolecular Energy (E_intermol)* included (1) the sum of components of Van der Waals interactions *(VdW),* the energy of Hydrogen bond formation *(H_bonds),* desolvation energy *(E_desolv)* and (2) Electrostatic energy (Electrostatic).

*Final Total Internal Energy* consists of interactions of atoms within the ligand and of interactions of atoms within the protein.

*Torsional Free Energy* is based on the difference of energies between the unbound and bound ligand, which is caused by the conformational changes depending on the number of rotatable bonds.

*Unbound System energy* is obtained for the ligand, which is supposed to be in an extended conformation in solution before ligand is bound to protein.

**Supplementary Figure legends**

**Figure S1. RITA binds to Np53 protein.**

A. Competition experiments show efficient displacement of p53/[^14^C]-RITA complexes, but not HSA/[^14^C]-RITA by unlabelled RITA (**B**).

**Figure S2. RITA analogs don’t interact with p53**

A. Model for the compound 3/p53 complex. SPLPS sequence of p53 is shown in cyan and NSC-672170 in magenta. Hydrogen bonds are highlighted in black dotted lines.
B. Model for compound 4 interaction with p53. SPLPS sequence of p53 is shown in cyan and NSC-650973 is indicated in green.

**Figure S3. RITA does not activate mouse p53 transcription activity.**

RITA and Nutlin efficiently induce p53 reporter in the ARN8 human melanoma cell line. Only Nutlin, not RITA induce p53 reporter in normal mouse fibroblasts.

**Figure S4. RITA induces compact conformation in wt Np53 but not in mutant Np53(33/37).**

A. nESI mass spectra (left) and collision cross-section distributions (CCSDs) (right) of 50 µM Np53 sprayed from 50 mM ammonium acetate in the absence of RITA (top panel), in the presence of RITA in a 1:2 ratio with 5% IPA (middle panel) and control spectra Np53 with 5% IPA (bottom panel). Charge state distributions are labeled with the lowest, highest and most intense charge state. Collision cross section distributions are derived from arrival time distributions at a drift voltage of 35 V for the [M+6H]^6+^ charge state of Np53. Conformational families are depicted as colored Gaussian curves. CCSDs are normalized to the intensity of each peak in the corresponding mass spectrum.
B. nESI mass spectra (left) and collision cross-section distributions (CCSDs) (right) of 50 µM Np53(33/37) sprayed from 50 mM ammonium acetate in the absence of RITA (top panel), in the presence of RITA in a 1:2 ratio with 5% IPA (middle panel) and control spectra Np53(33/37) with 5% IPA (bottom panel). Collision cross section distributions are derived from arrival time distributions at a drift voltage of 35 V for the [M+6H]^6+^ charge state of wt Np53. Conformational families are depicted as colored Gaussian curves. CCSDs are normalized to the intensity of each peak in the corresponding mass spectrum.

**Figure S5. RITA triggers conformational tightening at all charged states of wt Np53 but not of mutant Np53(33/37).**

A. Collision cross section distributions derived from arrival time distributions for wt Np53 in the absence (top panel) and presence (bottom panel) of RITA at charge states [M+5H]^5+^, [M+7H]^7+^, [M+8H]^8+^ and [M+9H]^9+^ all showing the conformational tightening of Np53 upon RITA incubation. CCSDs are normalized to the intensity of each peak in the corresponding mass spectrum.
B. Collision cross section distributions derived from arrival time distributions for Np53(33/37) in the presence and absence of RITA at charge states [M+5H]^5+^, [M+7H]^7+^ and [M+8H]^8+^ in the absence (top panel) and presence (bottom panel) of RITA. CCSDs are normalized to the intensity of each peak in the corresponding mass spectrum.

**Figure S6. Serine 33 and 37 are needed for the binding of repurposed protoporphyrin IX to p53 N-terminus.**

PpIX interaction with purified proteins, GST, GST-Np53, GST-Np53(33), GST-Np53(33/37) and human serum albumin (HSA), 20 μM incubated in 1:2 protein: PpIX molar ratio, was detected by fluorescent band shift assay as described previously (Sznarkowska et al., 2011). Band density was analyzed using ImageJ.

**Supplementary table 1. Structural analogs of RITA and their biological activities.**

**Movie S1. 3D animation of RITA/p53 complex**

A schematic 3D animation of the p53 N-terminal domain (in dark grey) bound to RITA (in blue). The N-terminal domain has been modeled as described in Methods and orientated with its N-terminus facing downwards. RITA-binding site (residues 32-37) is shown in teal (cyan) as on Figure 3. The MDM2-binding epitope (residues 19-26) is shown in pale green as on Figure 3. Residues of these two sites and RITA are shown in the sticks with nitrogen, oxygen, hydrogen and sulfur atoms shown in red, blue, light grey and yellow, respectively. Putative hydrogen bonds between RITA and serine residues 33 and 37 are shown as dotted lines. As evident from this model the RITA-bound state of the p53 N-terminal domain has the MDM2-binding residues (F^19^, W^23^, L^26^, in pale green) trapped inside of the hydrophobic cluster and inaccessible for MDM2 binding.
