## Supplementary figures and images for "Novel allosteric mechanism of dual p53/MDM2 and p53/MDM4 inhibition by a small molecule"

### Supplemental Figure 1

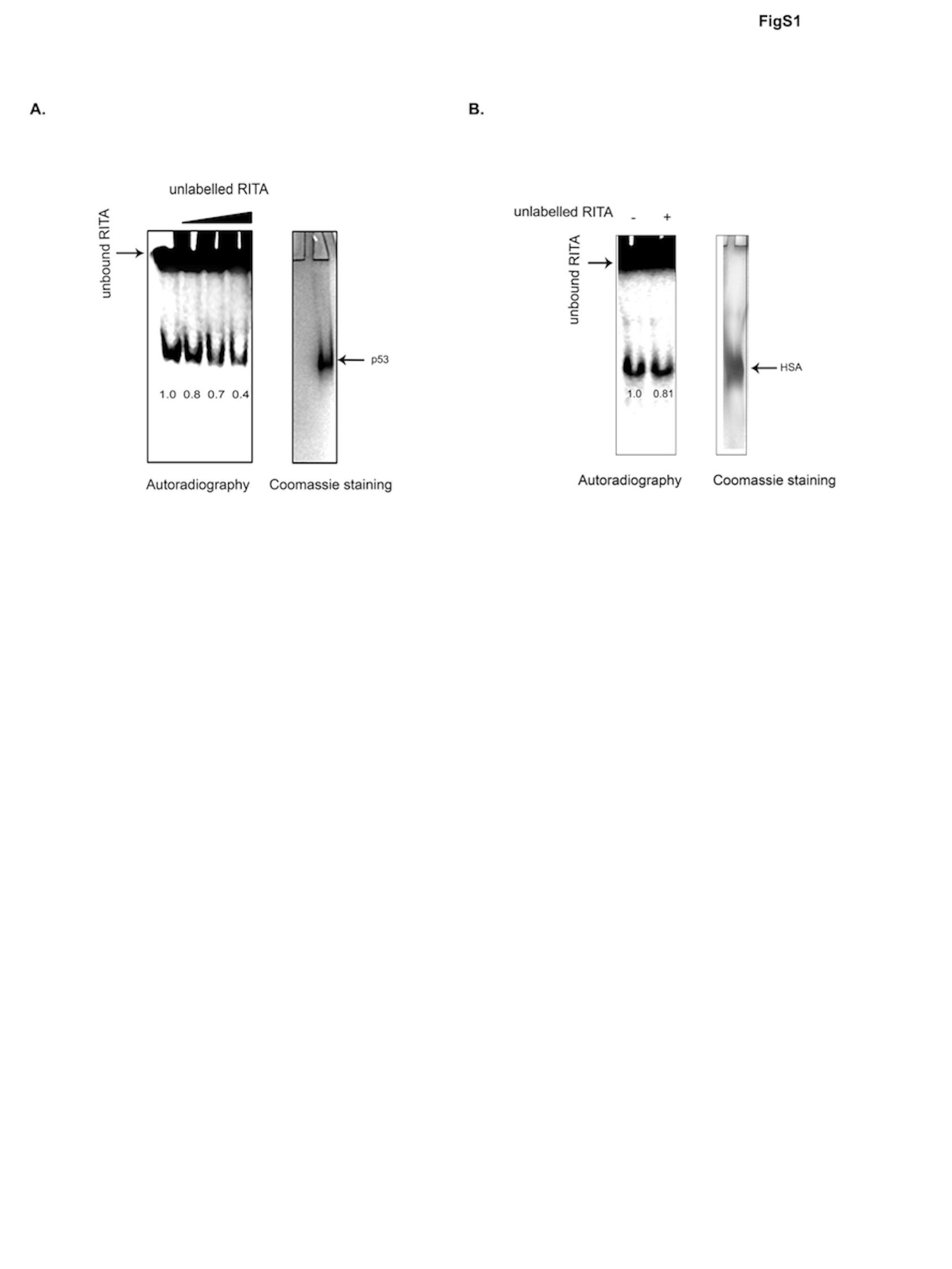

### Supplemental Figure 2

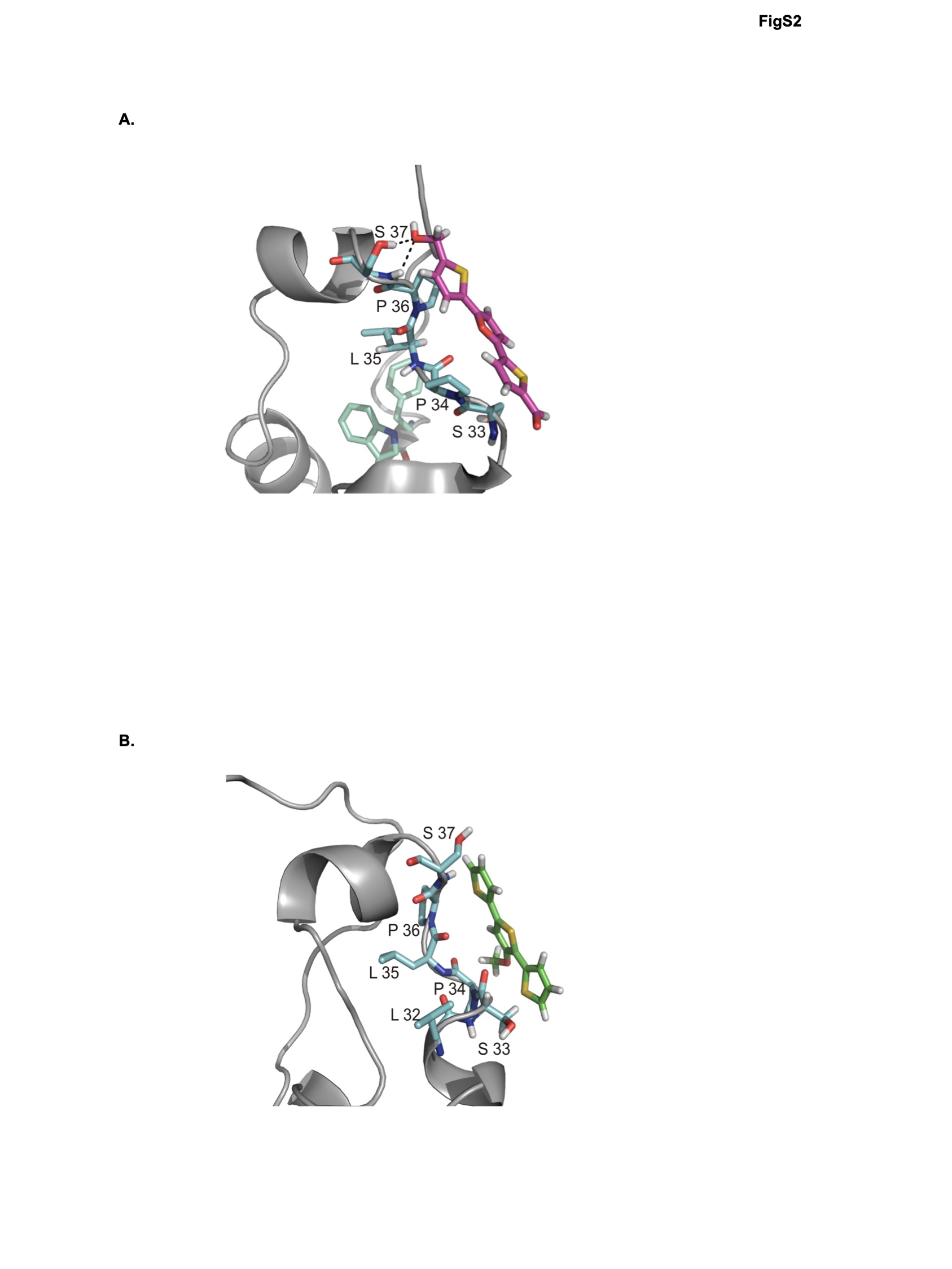

### Supplemental Figure 3

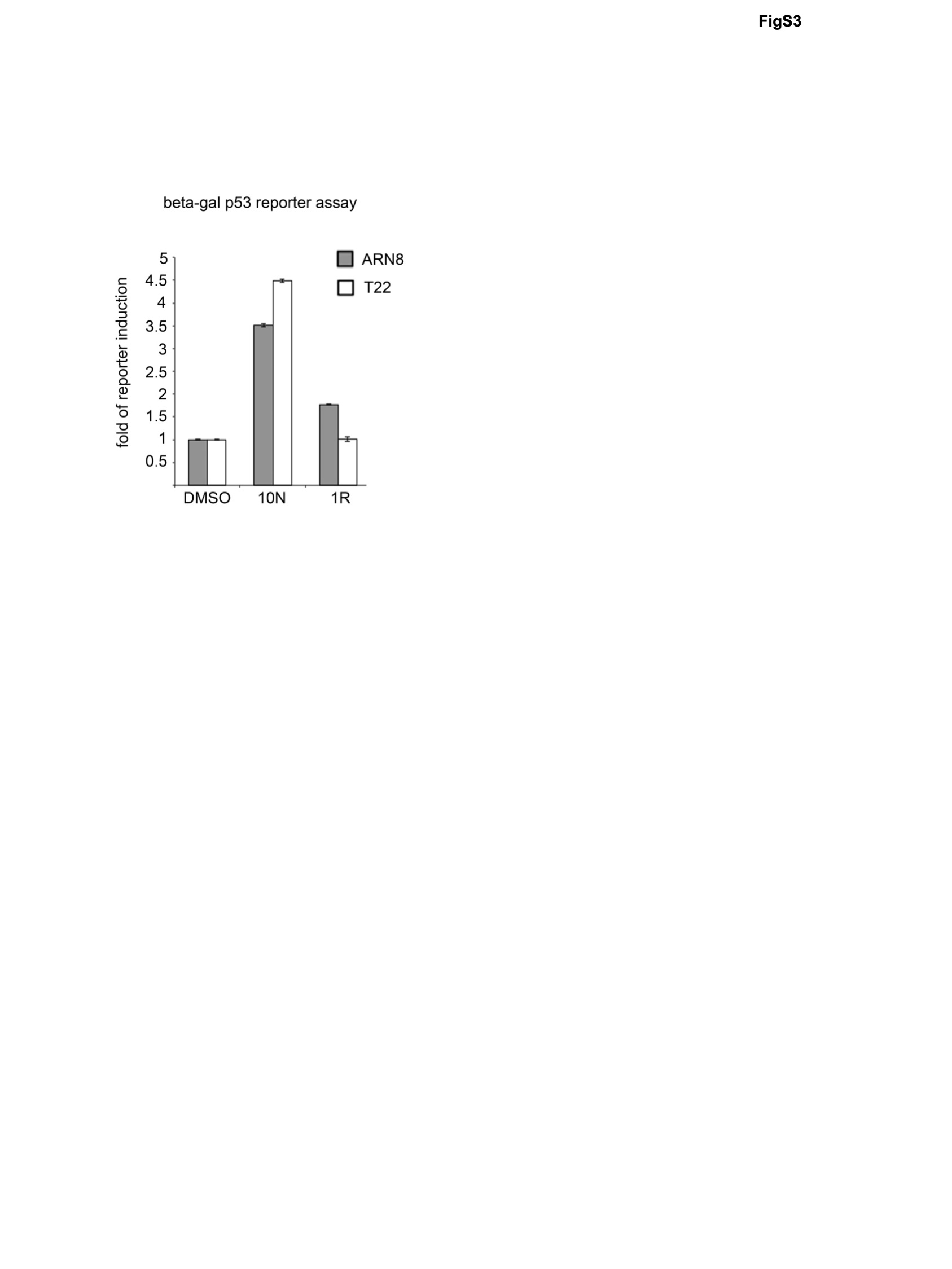

### Supplemental Figure 4

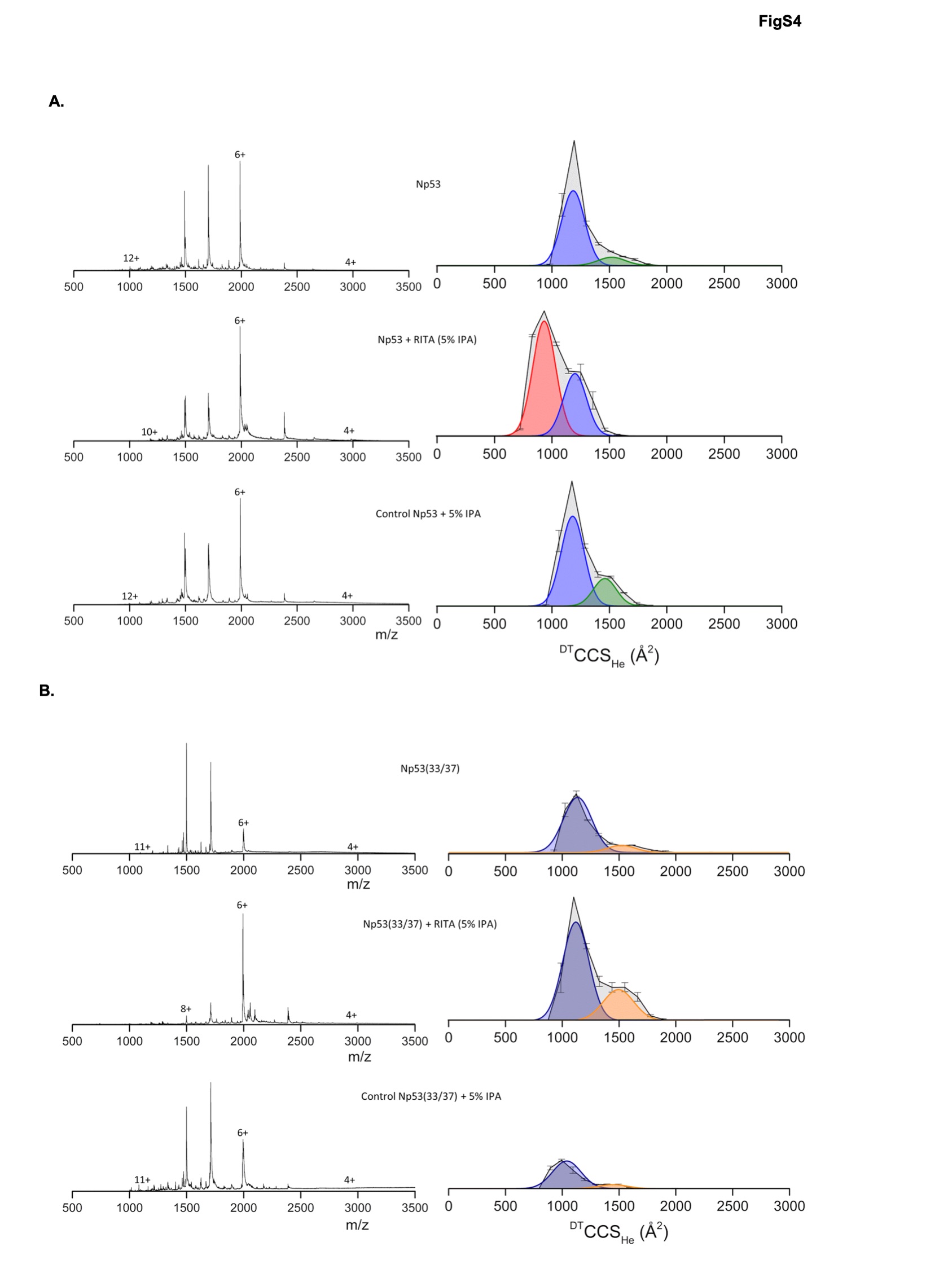

### Supplemental Figure 5

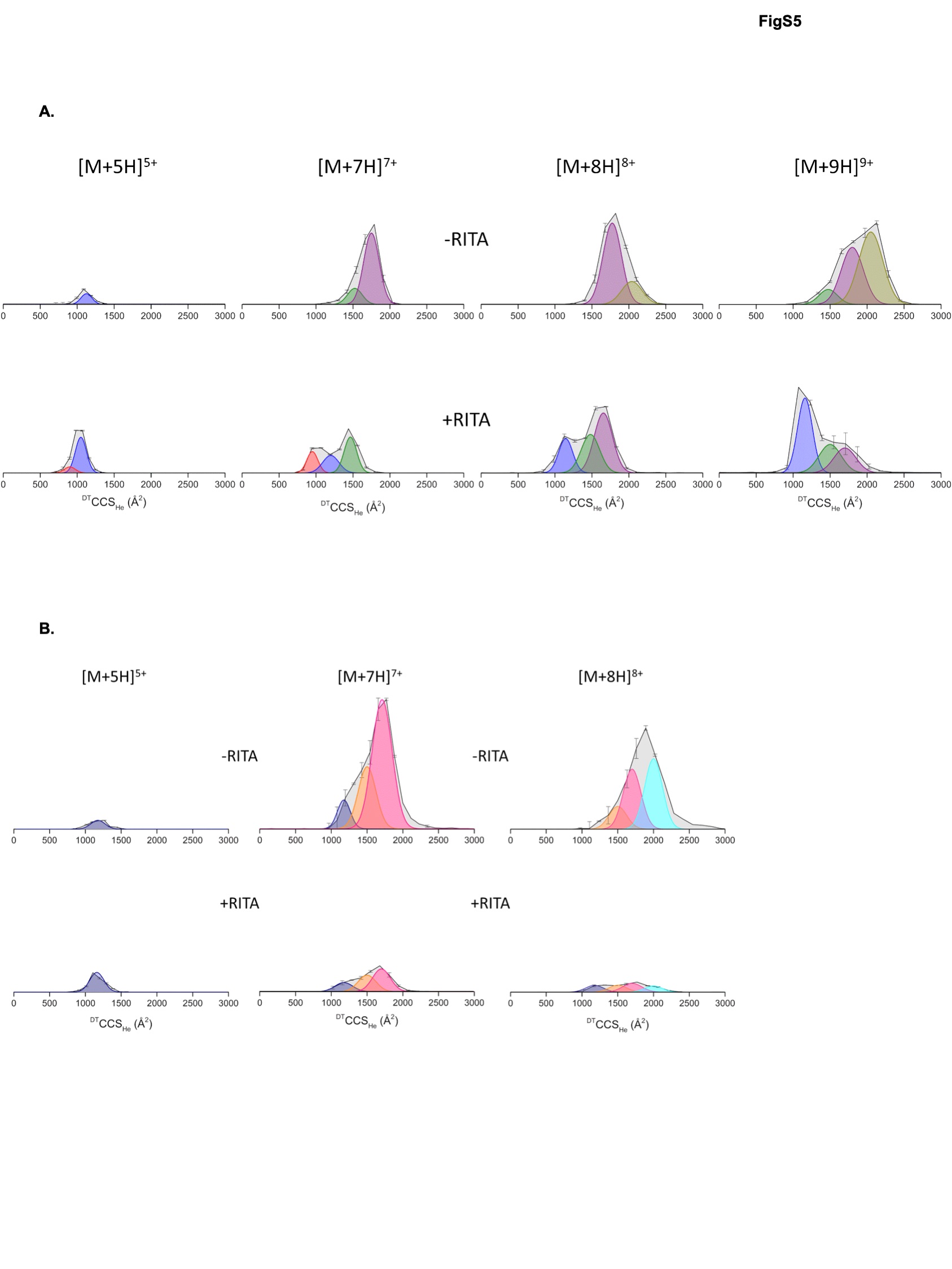

### Supplemental Figure 6

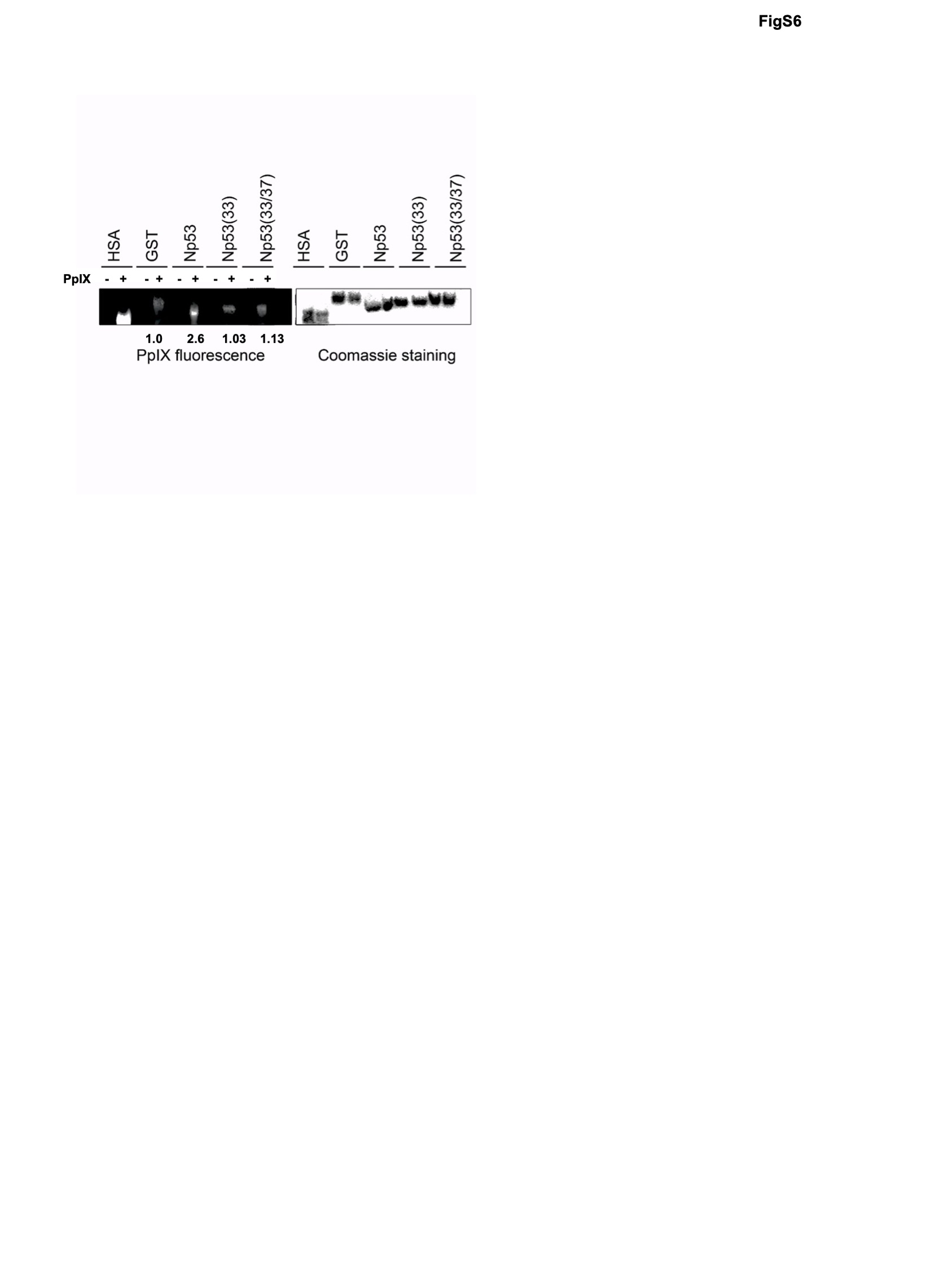

### Supplemental Table 1

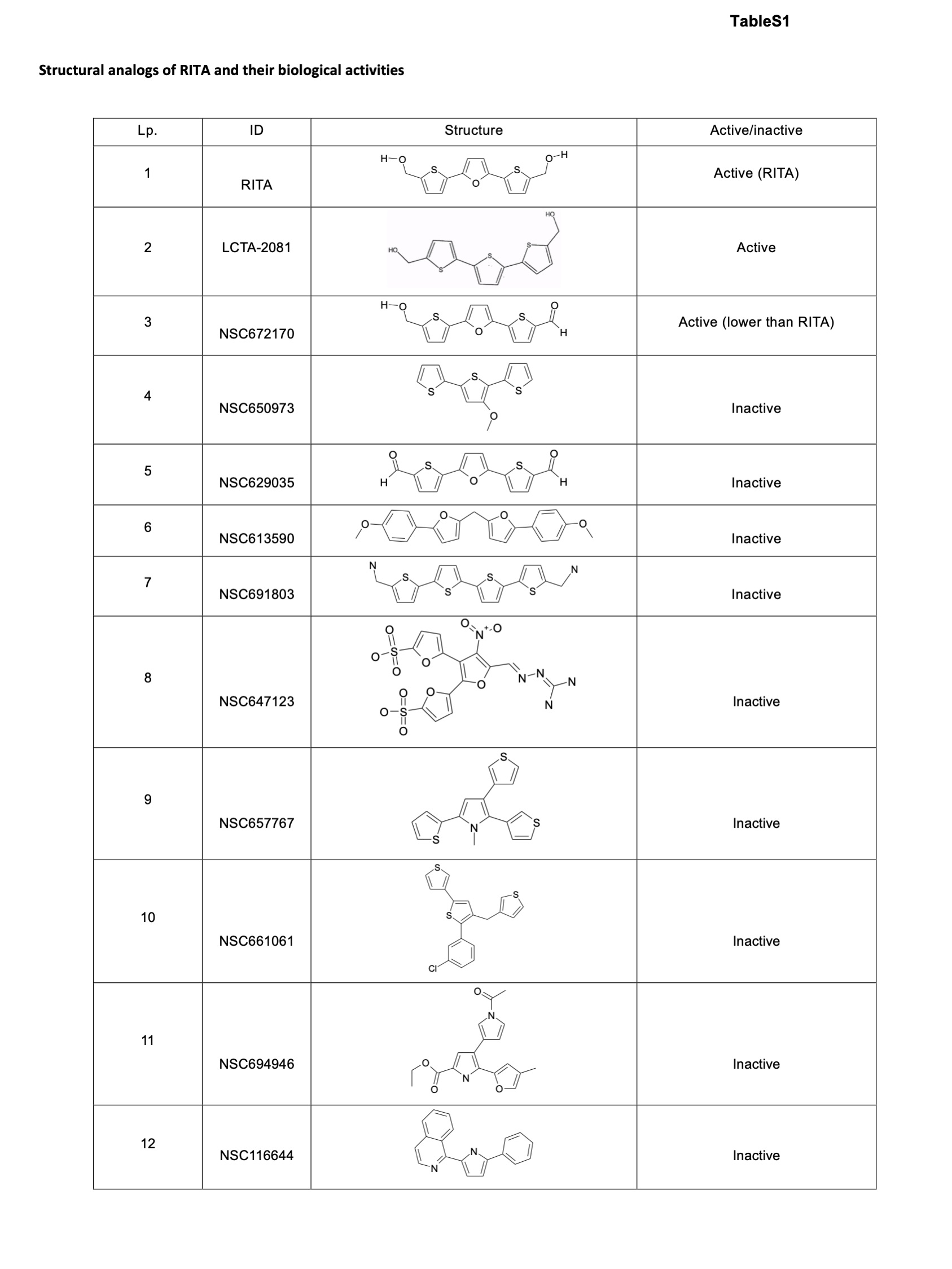
